# Mouse L1s fade with age: a methylation-enforced mechanism for attenuation of L1 retrotransposition potential

**DOI:** 10.1101/2022.08.06.500894

**Authors:** Patricia Gerdes, Dorothy Chan, Mischa Lundberg, Francisco J. Sanchez-Luque, Gabriela O. Bodea, Adam D. Ewing, Geoffrey J. Faulkner, Sandra R. Richardson

**Affiliations:** Mater Research Institute - University of Queensland, TRI Building, Woolloongabba QLD 4102, Australia.; The University of Queensland Diamantina Institute, The University of Queensland, Woolloongabba QLD 4102, Australia.; Translational Bioinformatics, Commonwealth Scientific and Industrial Research Organisation, Sydney NSW 2113, Australia.; GENYO. Centre for Genomics and Oncological Research (Pfizer-University of Granada-Andalusian Regional Government), PTS Granada, 18016, Spain.; MRC Human Genetics Unit, Institute of Genetics and Cancer (IGC), University of Edinburgh, Western General Hospital, Edinburgh EH4 2XU, United Kingdom.; Queensland Brain Institute, University of Queensland, Brisbane QLD 4072, Australia.

## Abstract

Mice harbor ∼2,800 intact copies of the retrotransposon Long Interspersed Element 1 (L1). The *in vivo* retrotransposition capacity of an L1 copy is defined by both its sequence integrity and epigenetic status, including DNA methylation of the monomeric units constituting young mouse L1 promoters. Locus-specific L1 methylation dynamics during development may therefore elucidate and explain spatiotemporal niches of endogenous retrotransposition, but remain unresolved. Here, we interrogate the retrotransposition efficiency and epigenetic fate of source (donor) L1s, identified as mobile *in vivo*. We demonstrate that promoter monomer loss consistently attenuates the relative retrotransposition potential of their offspring (daughter) L1 insertions. We also observe that most donor/daughter L1 pairs are efficiently methylated upon differentiation *in vivo* and *in vitro*. We employ Oxford Nanopore Technologies (ONT) long-read sequencing to resolve L1 methylation genome-wide and with locus-specific resolution, revealing a distinctive “smile” pattern in methylation levels across the L1 promoter region and thereby elucidating a molecular mechanism potentially underpinning L1 promoter shortening. Together, our results offer a novel perspective on the interplay between epigenetic repression, L1 evolution, and genome stability.

## INTRODUCTION

Retrotransposons are major contributors to ongoing mutagenesis in mammalian genomes. The autonomous non-LTR retrotransposon Long Interspersed Element 1 (LINE-1 or L1) is presently active in both humans and mice, and L1 sequences occupy ∼17% of human DNA and ∼18% of mouse DNA (Lander et al. 2001; Waterston et al. 2002). While humans contain a single active L1 subfamily, the mouse genome harbors three active L1 subfamilies, termed T_F_, G_F_, and A (Casavant and Hardies 1994; DeBerardinis et al. 1998; Goodier et al. 2001; Hardies et al. 2000; Mears and Hutchison 2001; Saxton and Martin 1998; Schichman et al. 1993; Wincker et al. 1987), which are each further divided into several sub-lineages, for example T_FI_, T_FII_ and T_FIII_ (Sookdeo et al. 2013). While T_F_ and G_F_ elements descended from the old and now inactive F subfamilies, A elements evolved independently. However, generating a single phylogenetic tree describing the relation of the subfamilies with each other is difficult due to frequent recombination among elements (Sookdeo et al. 2013). The vast majority of the ∼600,000 L1 copies in the mouse genome reference are 5ʹ truncated and mutated (Voliva et al. 1983; Waterston et al. 2002), leaving approximately 2,800 structurally intact L1s (Penzkofer et al. 2017). Ongoing L1 activity has generated substantial variation in L1 content among inbred strains, as well as inter-individual variation in L1 content within strains (Nellåker et al. 2012; Akagi et al. 2008; Richardson et al. 2017; Schauer et al. 2018). L1 insertions are also responsible for several spontaneous mouse mutants, driven by L1 T_F_ elements in all cases where the L1 subfamily can be identified (Gagnier et al. 2019).

A full-length mouse L1 is ∼6-7 kb long and begins with a 5ʹ untranslated region (5ʹ UTR) containing an internal RNA polymerase II promoter (DeBerardinis and Kazazian 1999; Severynse et al. 1991). The mouse L1 5ʹ UTR has a distinctive structure, wherein a variable number of tandemly repeated ∼200 bp monomer units are situated upstream of a non-monomeric region (Adey et al. 1994). Each monomer contributes additive promoter activity (DeBerardinis and Kazazian 1999). Individual monomers of young L1 subfamilies generally comprise sufficient CpG dinucleotides to qualify as CpG islands (Lee et al. 2010), and also contain several transcription factor binding sites, including for Yin Yang-1 (YY1) (Lee et al. 2010; DeBerardinis and Kazazian 1999). The YY1 binding site is required for accurate L1 transcription initiation (Lee et al. 2010; Athanikar et al. 2004). Consistent with YY1 dual functionality, an intact YY1 binding site is also important for methylation of the human L1 promoter during cellular differentiation (Sanchez-Luque et al. 2019).

The L1 5ʹ UTR is followed by two open reading frames encoding the proteins ORF1p and ORF2p, and a 3ʹ UTR incorporating a polyadenylation signal (Scott et al. 1987; Dombroski et al. 1991; Skowronski et al. 1988). ORF1p is ∼40kD and harbors RNA-binding and chaperone activities (Hohjoh and Singer 1996; Khazina and Weichenrieder 2009; Martin and Bushman 2001; Holmes et al. 1992; Khazina and Weichenrieder 2018) whereas the ∼150kD ORF2p has demonstrated endonuclease (EN) and reverse transcriptase (RT) activities (Doucet et al. 2010; Ergun et al. 2004; Feng et al. 1996; Mathias et al. 1991; Taylor et al. 2013). Both proteins are required for L1 mobilization through reverse transcription of an RNA intermediate in a process termed target-site primed reverse transcription (TPRT) (Luan et al. 1993; Feng et al. 1996; Holmes et al. 1992; Moran et al. 1996; Scott et al. 1987). While L1 retrotransposition can occur in non-dividing cells (Macia et al. 2017; Kubo et al. 2006), a growing body of evidence has emerged linking L1 retrotransposition to DNA replication during S-phase of the cell cycle (Flasch et al. 2019; Mita et al. 2018, 2020). Hallmarks of L1 integration by TPRT include flanking target site duplications (TSDs), and the incorporation of a 3ʹ poly-A tract which reflects the necessity of L1 mRNA polyadenylation for efficient retrotransposition (Grimaldi et al. 1984; Doucet et al. 2015).

Unchecked retrotransposition presents a threat to genome stability, and a variety of host defence mechanisms have evolved to curtail L1 activity (Goodier 2016; Greenberg and Bourc’his 2019; Bourc’his and Bestor 2004; MacLennan et al. 2017; Tristán-Ramos et al. 2020; Deniz et al. 2019; Senft and Macfarlan 2021; Liu et al. 2018; Mita et al. 2020). While repressive histone marks such as H3K9me3 play a major role in silencing older mouse L1 subfamilies (Castro-Diaz et al. 2014; Jacobs et al. 2014; Tan et al. 2013), younger L1s are typically silenced by methylation of the CpG islands in their promoters (Furano et al. 1988; Hata and Sakaki 1997; Lee et al. 2010; de la Rica et al. 2016). During embryonic development, the epigenome undergoes dramatic reprogramming including phases of global DNA demethylation (Seisenberger et al. 2012; Hajkova et al. 2002; Cantone and Fisher 2013; Saitou et al. 2012; Abe et al. 2011; Seki et al. 2005; Smith et al. 2012). The developmental methylation dynamics of L1 promoters have been examined using subfamily-specific and whole genome bisulfite sequencing (WGBS) (Saitou et al. 2012; Seisenberger et al. 2012; Smith et al. 2012; Hajkova et al. 2002; Popp et al. 2010; Watanabe et al. 2008; Kuramochi-Miyagawa et al. 2008; Molaro et al. 2014; Schöpp et al. 2020; Zoch et al. 2020). However, due to the repetitive structure and variable length of the mouse L1 promoter, as well as the high sequence identity among young L1 copies in the genome, assignment of short internal reads to specific mouse L1 loci is challenging (Lanciano and Cristofari 2020), and assessment of L1 methylation status *en masse* may mask individual L1s whose methylation dynamics differ to those of their subfamily. Indeed, studies of human L1s suggest certain loci can “escape” methylation and thus contribute to somatic retrotransposition throughout development and in cancer (Sanchez-Luque et al. 2019; Salvador-Palomeque et al. 2019; Nguyen et al. 2018; Schauer et al. 2018; Tubio et al. 2014; Philippe et al. 2016; Gardner et al. 2017; Paterson et al. 2015; Pitkänen et al. 2014; Ewing et al. 2020; Scott et al. 2016). Locus-specific resolution of murine L1 methylation, however, remains largely unexplored. Heritability of locus-specific retrotransposon methylation has been found in the context of “metastable epialleles” mostly involving variably methylated young IAP elements (VM-IAPs) (Bertozzi and Ferguson-Smith 2020). However, genome-wide screens have not revealed evidence of this phenomenon for L1s (Elmer et al. 2021; Kazachenka et al. 2018).

Here, we leverage the unique opportunity afforded by identification of source (donor) and their offspring (daughter) L1 germline insertions (Richardson et al. 2017) through uniquely L1 3ʹ transduced sequences (Xing et al. 2006; Goodier et al. 2000; Pickeral et al. 2000; Moran et al. 1999; Holmes et al. 1994; Beck et al. 2010), and access to the mosaic tissues from their originating animals and insertion-bearing offspring, to investigate the retrotransposition potential and epigenetic fate of *de novo* and active L1 copies *in vivo*. We also employ *in vitro* differentiation of mouse embryonic stem cells (mESCs) as a model to explore developmental L1 methylation dynamics at locus-specific resolution and genome-wide. By combining cell culture-based L1 retrotransposition assays, locus-specific bisulfite sequencing, and Oxford Nanopore Technologies (ONT) long-read DNA sequencing and methylation profiling, we uncover how developmental DNA methylation dynamics across the monomeric L1 promoter may play a key role in the ongoing conflict between L1 mutagenesis and the maintenance of germline genome stability.

## RESULTS

### L1 retrotransposition efficiency is diminished by ongoing promoter shortening

To evaluate the retrotransposition potential of *de novo* and polymorphic daughter elements relative to their donors, we PCR amplified, cloned and capillary sequenced all 5 donor/daughter pairs (Richardson et al. 2017) to derive the exact nucleotide sequence of each element (Sanchez-Luque et al. 2019) (see Materials and Methods). All 10 L1s contained at least one T_F_ monomer unit and encoded intact ORFs, and each daughter L1 contained between 0.6 and 1.8 fewer monomer units than the corresponding donor L1 (Table 1; Fig. 1A; Supplemental Fig. S1A). The remaining sequence of each daughter L1 was identical to its donor, with the exception of Insertion 2 which had a single nonsynonymous substitution in ORF1 (V303A) (Fig. 1A; Supplemental Fig. S1A).

**Figure 1.**
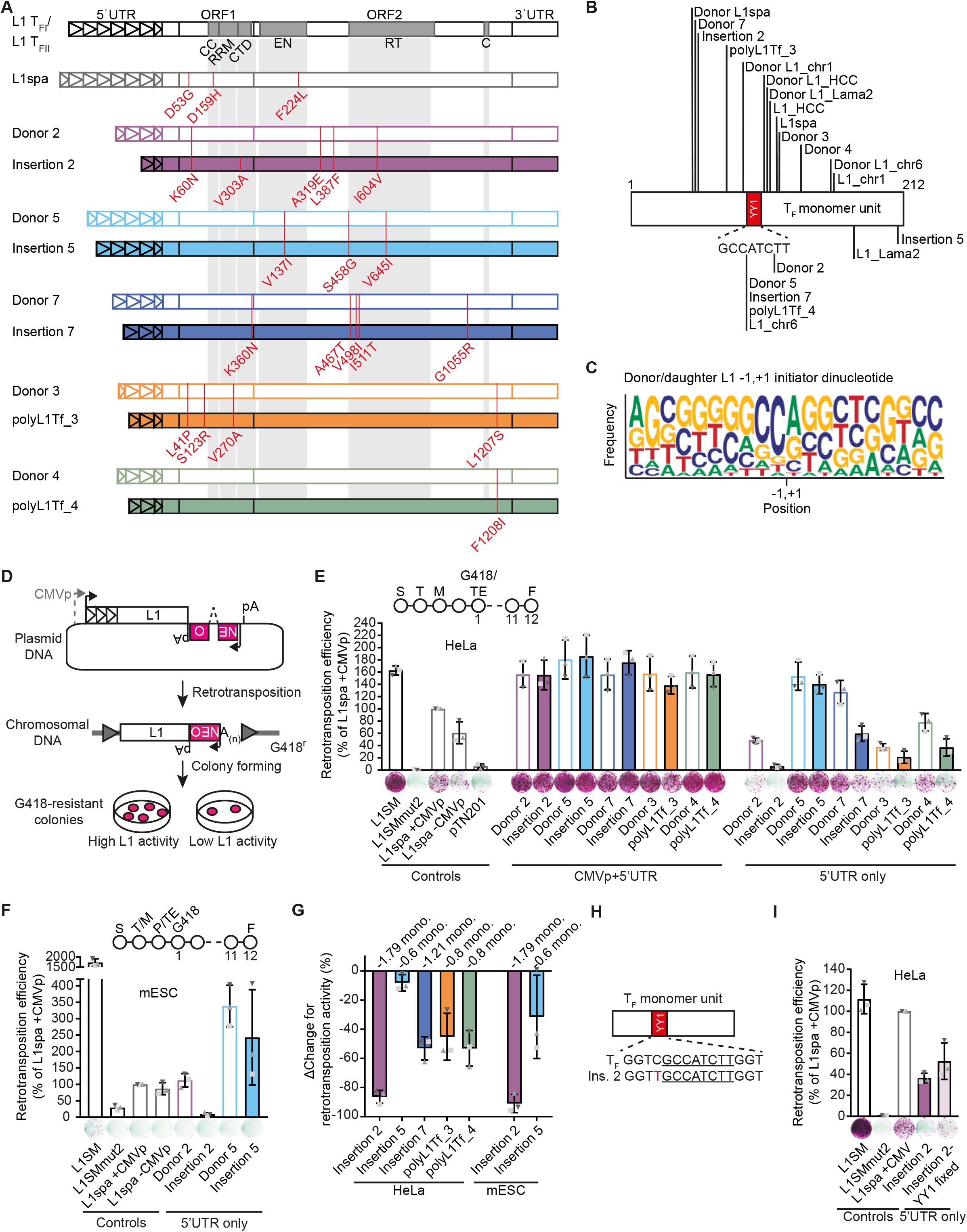
L1 donor/daughter pairs retrotranspose efficiently *in vitro*. (*A*) Amino acid changes in ORF1 and ORF2 compared to the L1 T_FI_ and L1 T_FII_ consensus sequences (Sookdeo et al. 2013) are annotated in red. Functional domains in ORF1 and ORF2 are shown: CC = coiled-coiled, RRM = RNA recognition motif, CTD = C-terminal domain, EN = endonuclease, RT = reverse transcriptase, C = cysteine-rich motif. Triangles within the 5ʹ UTR represent monomer units. For nucleotide substitutions in promoters see Supplemental Fig. S1. (*B*) Donor and daughter L1 5ʹ truncation points. Sequences or locations of donors and daughters were previously published as indicated in Table 1 (Richardson et al. 2017; Gagnier et al. 2019; Besse et al. 2003; Schauer et al. 2018; Kingsmore et al. 1994; Naas et al. 1998). Lines indicate truncation points of elements in the 5ʹ-most monomer. YY1 binding site (GCCATCTT) is shown in red. (*C*) Sequence logo (Crooks et al. 2004) of putative transcription initiation start sites for all donor and daughter L1s. Sequence represents transcription initiator dinucleotide in the center ± 9 nucleotides upstream and downstream. −1,+1 indicates transcription initiator dinucleotide. The first nucleotide of L1 sequence corresponds to the second nucleotide in transcription initiator dinucleotide. The position of the first base of each daughter L1 relative to its donor element, and the first base of each donor L1 relative to the L1 T_FI_/T_FII_ consensus sequences was analyzed. (*D*) Rationale of a cultured cell retrotransposition assay (Moran et al. 1996; Wei et al. 2000). Constructs used in this study were previously published (Moran et al. 1996; Goodier et al. 2001; Han and Boeke 2004) or generated by modifying the pTN201 construct (L1_spa_ (Naas et al. 1998)). An antisense orientated neomycin-resistance (NEO^r^) reporter cassette interrupted by a sense-oriented intron is inserted into a mouse L1 3ʹ UTR. The mouse L1 is driven by its native 5ʹ UTR promoter or a CMV promoter (CMVp). Cells harboring a retrotransposition event become neomycin (G418) resistant. The colony number reflects the relative activity of the L1 construct. (*E*) Comparison of L1 donor/daughter pair retrotransposition efficiency in HeLa cells. The retrotransposition assay timeline is shown in the *top* (S: seeding, T: transfection, M: change of media, G418: start of G418 selection, TE: measurement of transfection efficiency, F: Fixing and staining of colonies). Constructs: L1SM (positive control), L1SMmut2 (negative control), pTN201, L1_spa_ +CMVp/-CMVp, L1 donor/daughter pairs +CMVp/-CMVp. Colony counts were normalized to L1_spa_+CMVp and are shown as mean ± SD of three independent biological replicates, each of which comprised three technical replicates. Representative well pictures are shown below each construct. 5×10^3^ cells were plated per well in a six-well plate. (*F*) Comparison of L1 donor/daughter pair retrotransposition efficiency in mESCs. The retrotransposition assay timeline is shown at the *top* (S: seeding, T: transfection, M: change of media 8h after transfection, P: passaging of cells into 10 cm plates, TE: measurement of transfection efficiency, G418: start of G418 selection, F: Fixing and staining of colonies). Constructs as described in (*E*). Colony counts were normalized to L1_spa_+CMVp and are shown as mean ± SD of three independent biological replicates, each of which comprised two technical replicates. Representative well pictures are shown below each construct. 4×10^5^ cells were plated per well in a six-well plate. (*G*) Percentage change (ΔChange) in retrotransposition activity between L1 donor/daughter pairs. Shown is the decrease of retrotransposition efficiency per daughter L1 compared to its respective donor L1. Data is shown as mean ± SD of three independent biological replicates. (*H*) Schematic of an L1 monomer unit. The YY1 binding site is indicated as red rectangle. The extended YY1 binding motif sequence is shown below. The core YY1 binding motif sequence is underlined. A mutation in the extended YY1 binding motif sequence adjacent to the core motif in Insertion 2 is indicated in red. (*I*) Comparison of retrotransposition efficiency of Insertion 2 and Insertion 2 with intact YY1 binding sites (Insertion 2-YY1 fixed) in retrotransposition assay in HeLa cells. Constructs as described in (*D*). Colony counts were normalized to L1spa in pCEP4-mneoI-G4 + CMVp and are shown as mean ± SD of three independent biological replicates, each of which comprised three technical replicates. Representative well pictures are shown below each construct. 1×10^4^ cells were plated per well in a six-well plate. Note: L1SM retrotransposed very efficiently, leading to cell colony crowding in wells, and a likely underestimate of retrotransposition.

**Table 1.**
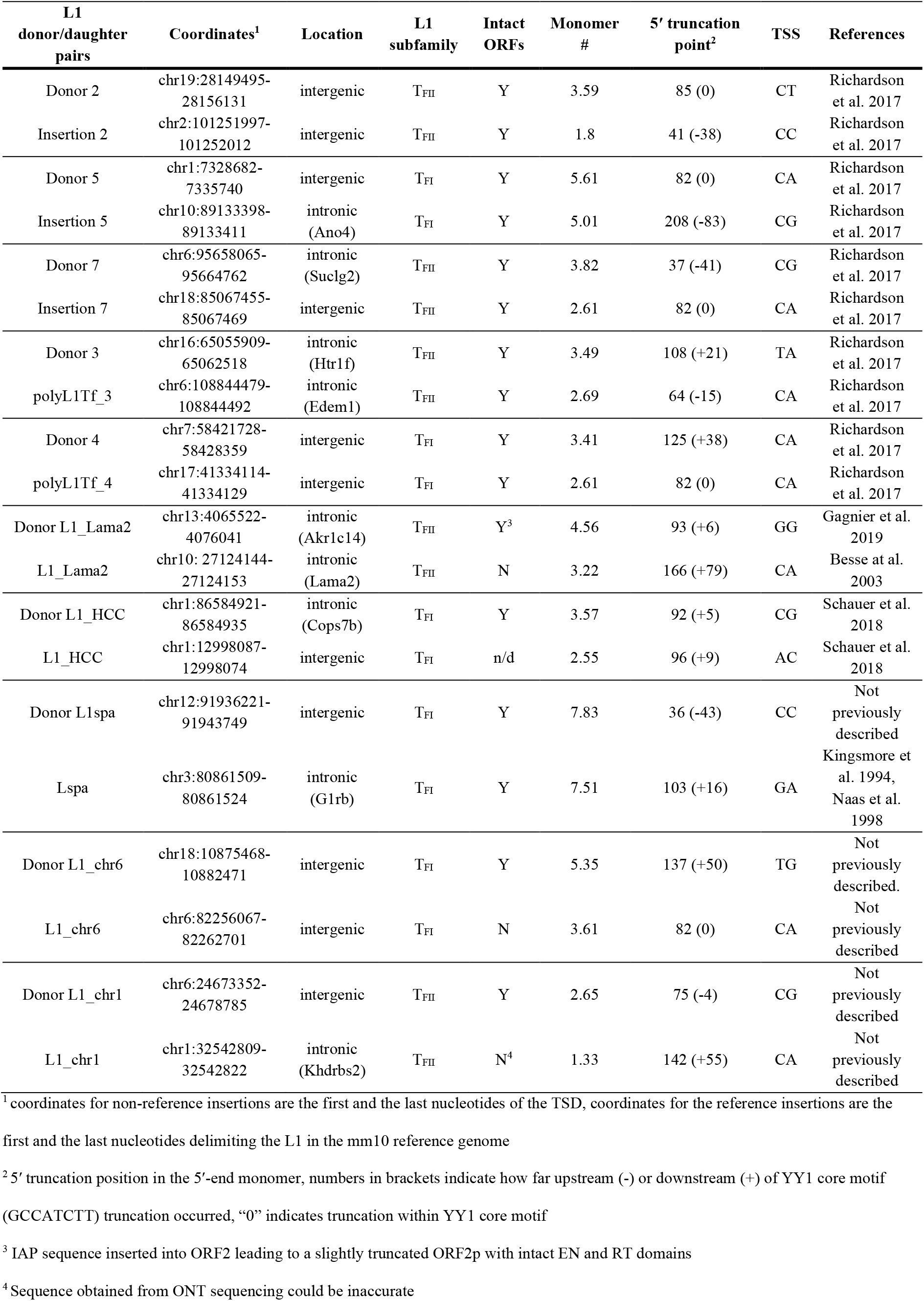
Characterization of L1 donor/daughter pairs.

The loss of daughter element 5ʹ UTR sequence could be explained either by 5ʹ truncation during retrotransposition (Ostertag et al. 2001; Symer et al. 2002; Zingler et al. 2005), or by the use of transcription start sites (TSSs) within internal monomers of the donor element promoter (DeBerardinis and Kazazian 1999). We analyzed the putative initiator dinucleotide (−1,+1) (Carninci et al. 2006) for the 10 elements under study (Fig. 1A), as well as 10 additional L1 T_F_ elements (Supplemental Fig. S1B) comprising 5 likely donor/daughter pairs (Table 1). Under the assumption that the most 5ʹ position of each element represents the first transcribed nucleotide, transcription of 14/20 elements initiated at the preferred mammalian PolII initiator pyrimidine/purine dinucleotide (Carninci et al. 2006) (Fig. 1C; Supplemental Fig. S1C; Table 1). 15 of the 20 elements analyzed here were 5ʹ truncated within the first 108 nt of the 5ʹ-most T_F_ monomer. 5/20 elements truncated within, and an additional 7/20 truncated in close proximity (≤21 bp) to, the YY1 core binding motif (GCCATCTT) (Fig. 1B; Table 1) as previously observed for mouse L1s (DeBerardinis and Kazazian 1999; Shehee et al. 1987; Zhou and Smith 2019). The observed clustering of 5ʹ truncation points, and their coincidence with the Py/Pu initiator dinucleotide, are consistent with the daughter L1 promoters being shortened due to transcription initiation internal to the 5ʹ UTR of the donor element.

To quantify the impact of monomer loss on daughter element mobility, we evaluated the 5 donor/daughter pairs identified in our previous study (Richardson et al. 2017) in a cultured cell L1 retrotransposition assay (Moran et al. 1996; Wei et al. 2000). We tested each element driven either by a cytomegalovirus promoter and the native L1 promoter (CMVp+5ʹ UTR), or by the native L1 promoter only (5ʹ UTR only), and quantified their activity relative to L1_spa_, a previously described disease-causing T_FI_ family insertion (Fig. 1D) (Naas et al. 1998; Kingsmore et al. 1994). In HeLa cells, when driven by CMVp+5ʹ UTR, all elements mobilized efficiently (∼160% of L1_spa_), with donor and daughter elements exhibiting similar activity. Notably, Insertion 2 which has an amino acid change in ORF1p retrotransposed with the same efficiency as its donor (∼160%) when transcribed from the CMVp, indicating that the mutation does not influence retrotransposition efficiency. In contrast, when driven by the 5ʹ UTR alone, each daughter element mobilized less efficiently than its donor (Fig. 1E,G). This trend was most pronounced for Insertion 2, which retrotransposed at 8% of L1_spa_+CMVp, compared to 62% for Donor 2 (Fig. 1E,G). A similar trend was observed when donor/daughter pairs 2 and 5 (5ʹ UTR only) were tested in mESCs (Fig. 1F,G) (MacLennan et al. 2017). We also noted that Insertion 2 had a single nucleotide mutation (1115C>T) in the extended YY1 binding motif (GGTCGCCATCTTGGT) found in its second monomer (Fig. 1H) which could increase YY1 binding affinity (Kim and Kim 2009). To determine whether the 1115C>T mutation contributes to the low retrotransposition efficiency of this element, we restored the YY1 binding site (Insertion 2-YY1 fixed), and observed a ∼15% increase in retrotransposition activity (Fig. 1I). Together, these retrotransposition assay results are consistent with previous luciferase reporter data showing mouse L1 promoter strength is proportionate to monomer number (DeBerardinis and Kazazian 1999). Moreover, we find that monomer loss consistently diminishes the retrotransposition potential of *de novo* mouse L1 insertions relative to their donor elements.

### Donor and daughter L1 insertions are largely methylated in adult tissues

Having established the retrotransposition competence of the donor and daughter elements *in vitro*, we next employed mouse L1 locus-specific bisulfite sequencing (Schauer et al. 2018) to evaluate the methylation status of each L1 in the somatic tissues and gonads of the mosaic animal in which the daughter insertion arose, and in subsequent generations of heterozygous insertion-bearing animals (Fig. 2A-D). For comparison, we analyzed the genome-wide methylation of the T_FI_ and T_FII_ subfamily monomers using primers internal to the monomer sequence (Schauer et al. 2018). Due to their high sequence similarity, it was not possible to design primers specific to the T_FI_ or T_FII_ subfamily. Overall, the T_FI_/T_FII_ subfamily monomer sequence was >80% methylated in adult tissues, although a few appreciably demethylated reads were observed across animals and tissues (Fig. 2E; Supplemental Fig. S2A).

**Figure 2.**
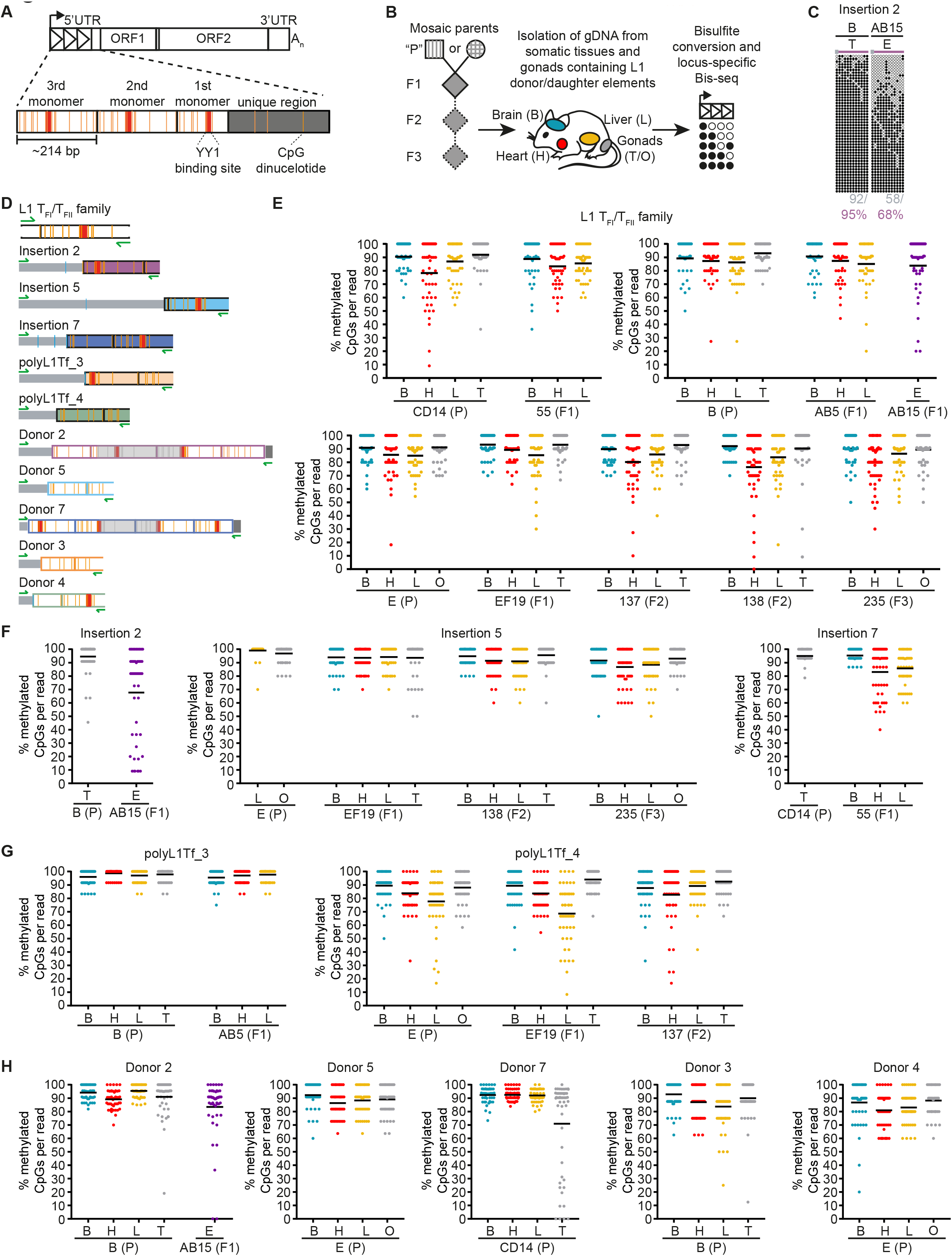
L1 donor/daughter elements are methylated in somatic tissues of adult mice. (*A*) Schematic of CpG dinucleotides in a mouse L1 T_F_ element. Triangles in 5ʹ UTR represent monomer units. A magnification of the 5ʹ UTR is shown below. Black boxes represent monomer units. Dark grey box represents unique (non-monomeric) region within 5ʹ UTR. Orange strokes represent CpG dinucleotides. Red boxes represent YY1 binding sites. (*B*) Experimental design of mouse L1 locus-specific bisulfite sequencing. Genomic DNA was extracted from tissues of C57BL6/J mice harboring previously identified donor and daughter L1 insertions (Richardson et al. 2017). The parental generation “P” (square = male, circle = female) is mosaic for the *de novo* daughter L1 insertion (represented by stripes). F1-F3 generations are heterozygous for L1 insertions (filled diamond = male or female). F2 and F3 generation animals were only available for Insertion 5 and polyL1Tf_4. Tissues for analysis of donor elements were only collected from the P generation. DNA was isolated from brain (green), heart (red), liver (orange) and gonads (grey) if available. After bisulfite conversion, the 5ʹ monomeric region of each L1 was PCR amplified. Amplicons were pooled and sequenced as 2×300mer Illumina reads. Circles represent methylated (black circles) and unmethylated (white circles) CpG dinucleotides in L1 5ʹ UTR. (*C*) Methylation of a *de novo* L1 promoter sequence (Insertion 2) shown in the germline mosaic P generation (animal B) and in the following F1 generation (animal AB15). Displayed are 50 non-identical sequences extracted at random from a much larger pool of available Illumina reads. Each cartoon panel corresponds to an amplicon (black circle, methylated CpG; white circle, unmethylated CpG; ×, mutated CpG). Colored line above cartoon represents amplicon (grey = genomic sequence, colored = L1 sequence). The overall percentage of methylated CpG dinucleotides is indicated below each cartoon. Grey letters indicate methylation of CpG dinucleotides in genomic sequence. Colored letters indicate methylation of CpG dinucleotides in L1 sequence. (*D*) Locus-specific methylation analysis schematic representation for L1 T_F_ monomer, 3 full-length *de novo* L1 insertions (Insertion 2, 5, 7), 2 polymorphic L1 insertions (polyL1Tf_3 and polyL1Tf_4) and their 5 respective donor elements (Donor 2, 5, 7, 3, 4). 5′ monomeric sequences of each L1 were PCR amplified using primer pairs (green arrows) specific to that locus. Orange strokes indicate L1 CpG dinucleotides covered by the assay. Blue strokes represent covered genomic CpG dinucleotides. Grey strokes in the grey shaded area represent CpG dinucleotides not reached by Illumina sequencing. Red boxes indicate YY1 binding sites. Colored boxes represent L1 monomer units. (*E*) Genome-wide methylation of L1 T_FI_/T_FII_ promoter sequence shown in all animals. Animals are labeled E, EF19, 137, 138, 235, B, AB5, AB15, CD14 and 55. The generation (P, F1, F2 or F3) is indicated in brackets. Each graph contains animals from the same family. Bisulfite sequencing was performed on DNA extracted from brain (B), heart (H), liver (L), testis (T) or ovaries (O) and embryonic tissues (E). Displayed are 50 non-identical sequences extracted at random from a much larger pool of available Illumina reads. Each dot in the diagram corresponds to an amplicon as per Supplemental Fig. S2A. The percentage of methylated CpGs per read is indicated on the *y*-axis. The average methylation is indicated by a black line. (*F*) As for (*E*) but for *de novo* L1 promoter sequences shown in the mosaic P generation where each *de novo* L1 insertion was identified (animals E and CD14) and in following F1-F3 generations if available (animals EF19, 138, 235 and 55). Only CpG dinucleotides within L1 sequences (represented as orange strokes in (*D*)) were analyzed. The average methylation of each promoter is indicated by a black line. (*G*) As for (*E-F*) but for polymorphic L1 insertions. (*H*) As for (*E-F*) but showing methylation of 5 donor L1 elements in the animal they mobilized in (P generation).

The three *de novo* daughter L1s (Insertions 2, 5, and 7) (Table 1) were >80% methylated in the somatic tissues and gonads of adult mice, including mosaic animals E (Insertion 5), CD14 (Insertion 7), and their heterozygous F1, F2, and F3 descendants (Fig. 2C,F; Supplemental Fig. S2B). Notably, Insertion 2 originated as germline-restricted mosaic in mouse B and was transmitted only to F1 animal AB15, which was harvested as a post-implantation embryo. Insertion 2 was highly methylated in the adult testis of mouse B, but partially demethylated in embryo AB15 (Fig. 2C,F). As the genomic DNA of embryo AB15 was analyzed in bulk, the demethylated sequences potentially corresponded to PGCs or multipotent stem cells. Consistently, the T_FI_/T_FII_ subfamily monomer sequence and Donor 2 (see below) also exhibited partial demethylation in embryo AB15. We also analyzed the methylation status of unfixed polymorphic insertions polyL1Tf_3 and polyL1Tf_4 (Table 1). Insertion polyL1Tf_3 was nearly 100% methylated across all adult tissues examined (Fig. 2G; Supplemental Fig. S2B). However, methylation of polyL1Tf_4 was more relaxed (<90%), with a tendency to be especially demethylated in liver (Fig. 2G; Supplemental Fig. S2B). This variability may reflect the influence of genomic location and physiological context on L1 element methylation (Sanchez-Luque et al. 2019; Salvador-Palomeque et al. 2019; Ewing et al. 2020). Together, these results indicate that *de novo* L1 insertions arising during embryonic development are likely silenced by DNA methylation during later embryogenesis and this methylation is maintained in subsequent generations with an average methylation level of 93% in brain, 89% in heart and 89% in liver for daughter insertions and 92% in brain, 87% in heart and 87% in liver for the donor L1s.

We viewed donor L1s active during embryonic development as candidate “escapee” loci in mice, potentially akin to specific human L1 loci that evade epigenetic repression in differentiated cells (Sanchez-Luque et al. 2019; Salvador-Palomeque et al. 2019; Ewing et al. 2020; Scott et al. 2016). We therefore assessed methylation of Donor 2, Donor 5 and Donor 7 in somatic tissues and germ cells of their mosaic founder animals (mouse B, mouse E, and mouse CD14, respectively). We also analyzed methylation of Donor 3 and Donor 4 in animals that carried the respective polymorphic daughter insertions polyL1Tf_3 and polyL1Tf_4 (Fig. 2B,D). Nearly all donor elements showed >80% methylation in somatic tissues and gonads (Fig. 2H; Supplemental Fig. S2C). The exception was Donor 7 which, while completely methylated in the somatic tissues of founder mouse CD14, was hypomethylated in the germ cell fraction of mouse CD14 testis. This is a notable departure from the genome-wide state of T_FI_/T_FII_ monomer sequences, which we found to be largely methylated in adult gonads (Fig. 2E,H; Supplemental Fig. S2A,C). Thus, Donor 7 may represent an L1 that is refractory to methylation during germline development and therefore privileged for heritable retrotransposition, while the majority of mouse L1s actively mobilizing during embryonic development are efficiently methylated in adult tissues.

### Young L1s are rapidly methylated during *in vitro* mESC differentiation

As adult tissues reveal only the end-point of developmental L1 methylation dynamics, we next analyzed L1 methylation at genome-wide and locus-specific resolution during cellular differentiation. To model various states of pluripotency, we cultured feeder-free E14 mouse embryonic stem cells (mESCs) in three conditions: serum complemented with leukemia inhibitory factor (serum+LIF), which generates a heterogeneous population of mESCs in a pluripotent state (Smith et al. 1988; Williams et al. 1988); two small kinase inhibitors + LIF (2i+LIF), which maintains the mESCs in a naïve ground state similar to that of the inner cell mass (ICM) (Silva et al. 2008); and under 2i+serum conditions, which are shown to support engineered mouse L1 retrotransposition (MacLennan et al. 2017). To recapitulate specification and differentiation of cells into the three germ lineages (ectoderm, mesoderm and endoderm), we collected genomic DNA over a time-course, from differentiation induction of serum+LIF mESCs to embryoid bodies (EBs) for 6 days, and through subsequent differentiation over 15 days (Fig. 3A-B; Supplemental Fig. S3A-B) (Behringer et al. 2016). L1 methylation state was assessed using genome-wide and locus-specific L1 bisulfite sequencing in the three mESC culture conditions and on days 3, 6, 9, 12, 15, 18 and 21 of differentiation.

**Figure 3.**
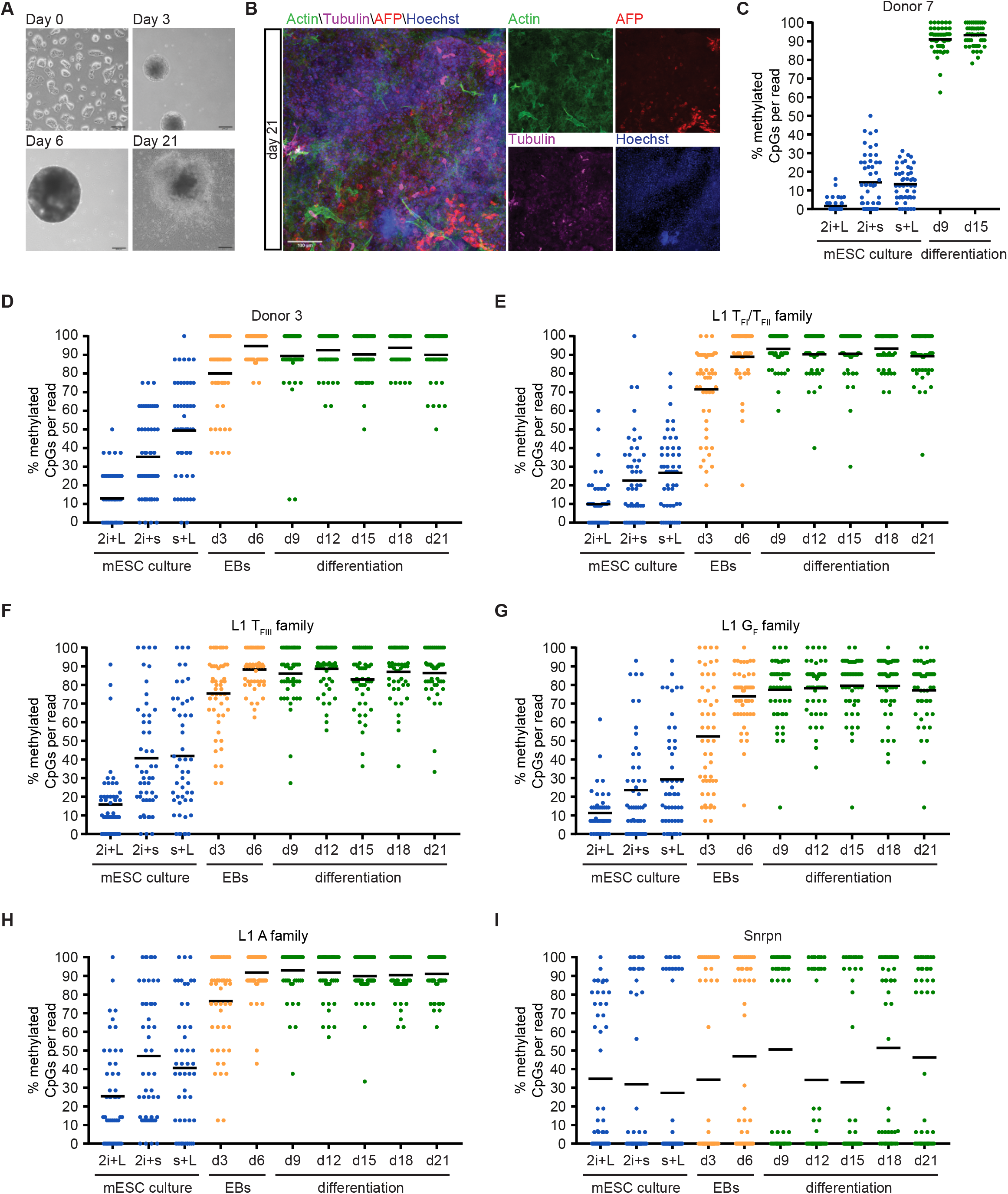
Dynamic methylation of L1 elements during differentiation of mESCs. (*A*) Differentiation of mESCs to cells of all three germ layers using a standard differentiation protocol (Behringer et al. 2016). Undifferentiated E14 mESCs are grown on gelatin (day 0). Embryoid bodies (EBs) are generated by “hanging drop culture” (day 3) and are grown in suspension culture (day 6). After six days, EBs are plated and differentiated for two weeks (day 21). Scale bar, 200 µm. (*B*) Immunofluorescence image of mesodermal (Actin, green), endodermal (AFP, red) and ectodermal (Tubulin, violet) lineage markers in differentiated E14 mESCs on day 21. Nuclei were stained with Hoechst (blue). Scale bar, 100 µm. (*C*) Methylation of Donor 7 promoter sequence shown in the mESCs cultured in three different conditions (2i+L = 2i+LIF, 2i+s = 2i+serum, s+L = serum+LIF) and on differentiation day 9 (d9) and day 15 (d15). Due to the technical challenge posed by PCR amplification of long bisulfite-treated fragments, sufficient material was generated to assess Donor 7 methylation only at day 9 and day 15 of differentiation. Displayed are 50 non-identical sequences extracted at random from a much larger pool of available Illumina reads. Each dot in the diagram corresponds to an amplicon. The percentage of unmethylated CpGs per read is indicated on the *y*-axis. Only CpG dinucleotides within L1 sequences represented as orange strokes in Fig. 2D were analyzed. The average methylation of each promoter is indicated by a black line. (*D-I*) As for (*C*) but for Donor 3 (*D*), L1 T_FI_/ T_FII_ family (*E*), L1 T_FIII_ family (*F*), L1 G_F_ family (*G*), L1 A family and the imprinted gene *Snrpn* (*I*). Primers for L1 families are within the L1 promoter sequence. Shown is methylation in three different mESC culture conditions, during EB culture and during differentiation.

The mobile L1 subfamilies T_F_, G_F_ and A each exhibited their lowest methylation levels in 2i+LIF ground-state conditions, and reached maximal methylation by day 6 of differentiation (Fig. 3E-H; Supplemental Fig. S4C-F). Notably, the G_F_ subfamily was less methylated (<80%) than the other subfamilies (>80%) both during differentiation and in fully differentiated EBs (Fig. 3G; Supplemental Fig. S4F). The 129/Ola genetic background from which E14 mESCs are derived contained two of the donor L1s analyzed above, Donor 3 and Donor 7. Both loci were largely demethylated (<50%) across all mESC culture conditions (Fig. 3C-D; Supplemental Fig. S4A-B). Donor 3 showed 80% methylation at day 3 of EB differentiation, and >90% at day 6. Donor 7 showed >90% methylation at these time points (Fig. 3C; Supplemental Fig. S4A). These results were in line with our observation that both donors were completely methylated in somatic tissues of adult mice (Fig. 2H) and is consistent with methylation occurring during differentiation in embryonic development *in vivo*. As an internal control, analysis of the maternally imprinted *Snrpn* gene revealed the expected bimodal distribution of methylation in pluripotent cells and across the differentiation time course (Fig. 3I; Supplemental Fig. S4G). Together, these results demonstrate rapid remethylation of L1 sequences during cellular differentiation, with subtle but notable variability between active L1 subfamilies and among individual loci.

### Methylation fluctuates within mouse L1 promoters

Although our locus-specific bisulfite sequencing approach allows base-pair resolution of individual L1 methylation, it is limited to the 5ʹ-most portion of each L1 promoter. To attain complete methylation profiles of L1 loci without bisulfite conversion, we performed PCR-free ONT sequencing of mESCs in serum+LIF (d0), d3 EBs, and d21 differentiated cells. We achieved ∼15-20× genome-wide depth using an ONT PromethION platform, and surveyed CpG methylation via the methylartist package (Ewing et al. 2020; Cheetham et al. 2021). DNA methylation evaluated by ONT sequencing increased during differentiation (Fig. 4A). Examining the methylation profiles of each L1 subfamily averaged across their ∼7kb consensus sequences, we found that in d0 mESCs full-length T_F_, G_F_ and A subfamily L1s were less methylated compared to the older L1 F subfamily (Fig. 4A). Average methylation of all L1 subfamilies rapidly increased to ∼80% by day 3 of differentiation; however, T_FI_ and T_FII_ elements were less methylated (≤80%) in d3 EBs compared to other L1 subfamilies (Fig. 4A). B1, B2 and BC1 SINEs were only slightly demethylated in d0 mESCs (∼80%), and fully methylated (>90%) in d3 EBs and d21 fully differentiated cells (Fig. 4B). While IAP retrotransposon LTRs were generally almost 100% methylated at all three differentiation timepoints, the IAP subfamily IAPEY4 LTRs showed an average methylation level of <40% in d0 mESCs and only∼70% methylation in d3 EBs and d21 differentiated cells. MERVL/MT2 LTRs were >80% methylated at all three timepoints. Etn elements showed an average methylation level of only ∼60% in d0 mESCs but >80% in d3 EBs and d21 differentiated cells (Fig. 4C).

**Figure 4.**
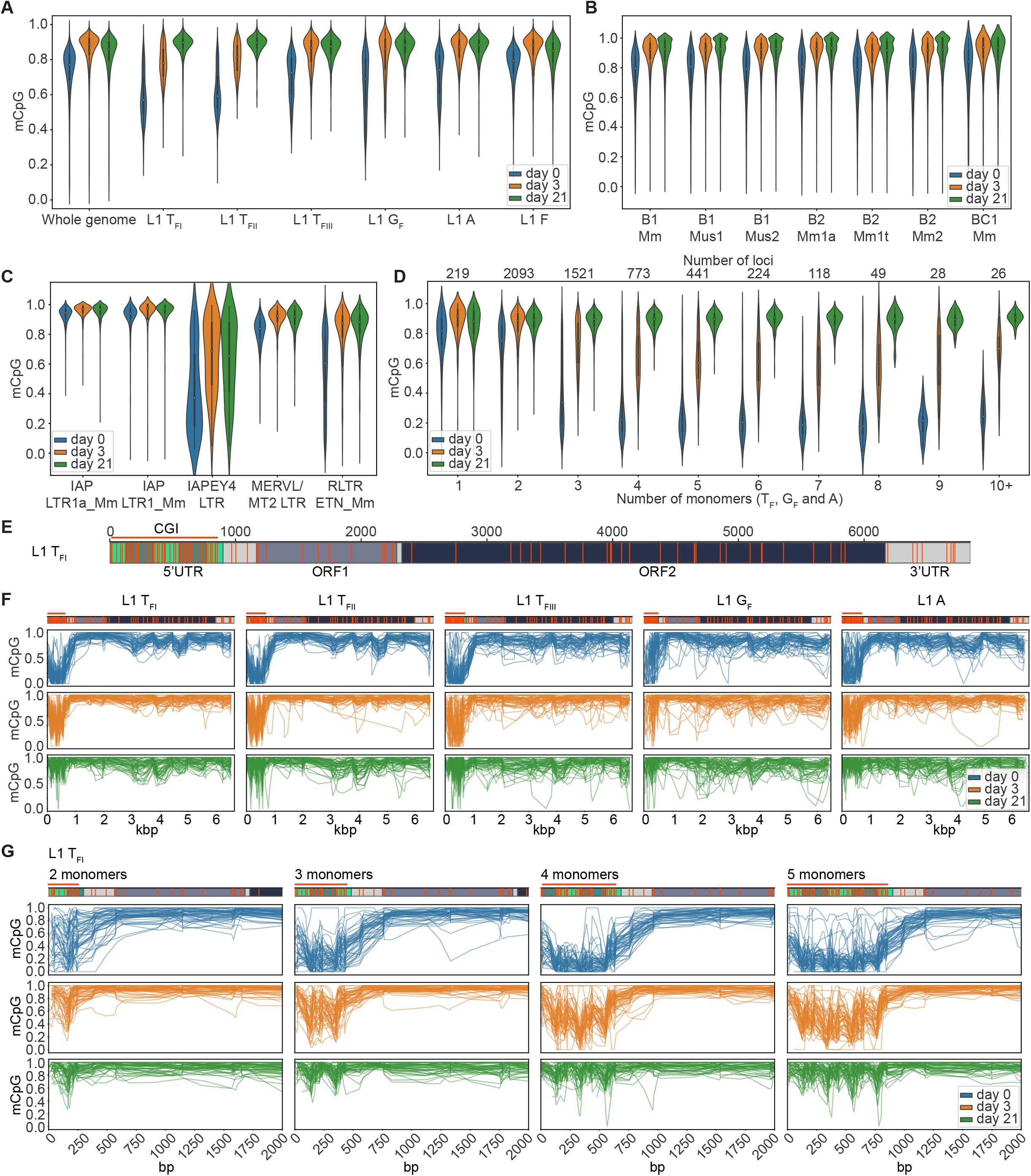
ONT CpG methylation profiles of TEs. (*A*) Violin plots are showing methylated CpG fraction for the whole genome (6 kbp windows), entire L1s belonging to the active L1 T_FI_, T_FII_, T_FIII_, G_F_ and A subfamilies and the evolutionary older and inactive L1 F subfamily at three time points of differentiation: d0 (undifferentiated mESCs in serum+LIF), d3 (EBs on day 3 of differentiation) and on d21 (completely differentiated cells). (*B*) As for (*A*), but for B1, B2 and BC1 SINE subfamilies. (*C*) As for (*A*), but for IAP LTR1a_Mm and LTR1_Mm, IAPEY4 LTR, MERV-L/MT2 LTR and RLTR ETn_Mm copies. (*D*) As for (*A*), but for violin plots showing the methylated CpG fraction of active L1 subfamily (T_F_, G_F_ and A together) promoters (monomers only) depending on the number of monomers. Only elements with a minimum coverage of 5 reads across the whole 5ʹ UTR were included in the plot. The number of loci represented in each bin is shown in the *top*. (*E*) Annotated full-length L1 T_F1_ consensus showing the monomer units in green, unique region in light grey, ORF1 in dark grey, ORF2 in dark green, and 3ʹ UTR in light grey. CpG dinucleotides throughout the whole element are displayed as orange strokes. The promoter CpG island (CGI) is indicated as an orange line. Number of bp are shown above the element. (*F*) Data is shown for full-length L1s (T_FI_, T_FII_, T_FIII_, G_F_ and A) containing 4 monomers in the promoter at three time points of differentiation: d0 (undifferentiated mESCs in serum+LIF), d3 (EBs on day 3 of differentiation) and on d21 (completely differentiated cells). Each graph displays up to 50 methylation profiles for the specified L1 subfamily. Annotated consensus sequences as per (*E*) are shown at *top* including CpG positions. (*G*) As for (*F*), but for promoters of L1 T_FI_ subfamily members containing between 2 and 5 monomers at three time points of differentiation (d0, d3 and d21).

Next, we quantified the methylation of active L1 subfamily promoters independently of the L1 body (unique 5ʹ UTR region, ORFs, 3ʹ UTR). We binned promoters genome-wide based on monomer count, with the majority of young L1 subfamily members containing between 2 and 5 monomer units (Fig. 4D), consistent with previous analyses of the mouse reference genome (Zhou and Smith 2019). On the whole, L1 promoter methylation was distributed between ∼0% and 90% in the d0 mESCs, indicating variability among loci even in pluripotent serum+LIF culture conditions. Methylation of L1 loci in d3 EBs was higher than in d0 mESCs but a substantial proportion of L1s were still <80% methylated. Notably, none of the L1 loci displayed here appeared to be completely demethylated in d3 EBs. Methylation of the majority of L1s was re-established (>80%) in d21 differentiated cells (Fig. 4D). However, some loci appeared to be <70% methylated even in d21 differentiated cells. We examined three of these methylation “escapees” and found that each of them showed a mixture of methylated and demethylated reads at d21 potentially belonging to specific cell types in our mixed population of differentiated cells (Supplemental Fig. S5) indicating again a likely lineage or cell-type specificity for L1 methylation “escapees” (Sanchez-Luque et al. 2019; Salvador-Palomeque et al. 2019; Ewing et al. 2020). Based on the mouse genome reference sequence all three L1s contain intact ORFs and multiple (4-7) monomers, indicating their potential retrotransposition competence.

We next assessed composite methylation profiles covering the previously inaccessible interiors of full-length mouse L1s and in particular the entire mouse L1 promoter (Fig. 4E). We observed a consistent methylation trough in the 5ʹ UTR promoter region in d0 mESCs and d3 EBs, whereas the L1 body was consistently hypermethylated at all three timepoints (Fig. 4F). Zooming in on the promoter region of the composite L1 methylation profiles revealed a consistent “smile” pattern across the 5ʹ UTR, with the innermost monomers less methylated compared to the 5ʹ-most and 3ʹ-most monomers (Fig. 4G; Supplemental Fig. S6). This methylation pattern was most pronounced in T_FI_ and T_FII_ elements (Fig. 4G; Supplemental Fig. S6A-D). Our analyses also revealed peaks and valleys of methylation along mouse L1 promoters, with a periodicity corresponding to the monomer units. Taking advantage of the locus-specific resolution offered by ONT sequencing, we examined methylation of the two donor L1s (Donor 3 and Donor 7) present in E14 mESCs (Fig. 5). These elements recapitulated the methylation trough observed in the composite plots in d0 mESCs and d3 EBs. As an internal control, we readily identified the differentially methylated regions (DMRs) of two imprinted genes, *Snrpn* and *Impact* (Supplemental Fig. S7). In sum, our ONT methylation analysis of full-length L1s provides unprecedented resolution of interior mouse promoter methylation dynamics.

**Figure 5.**
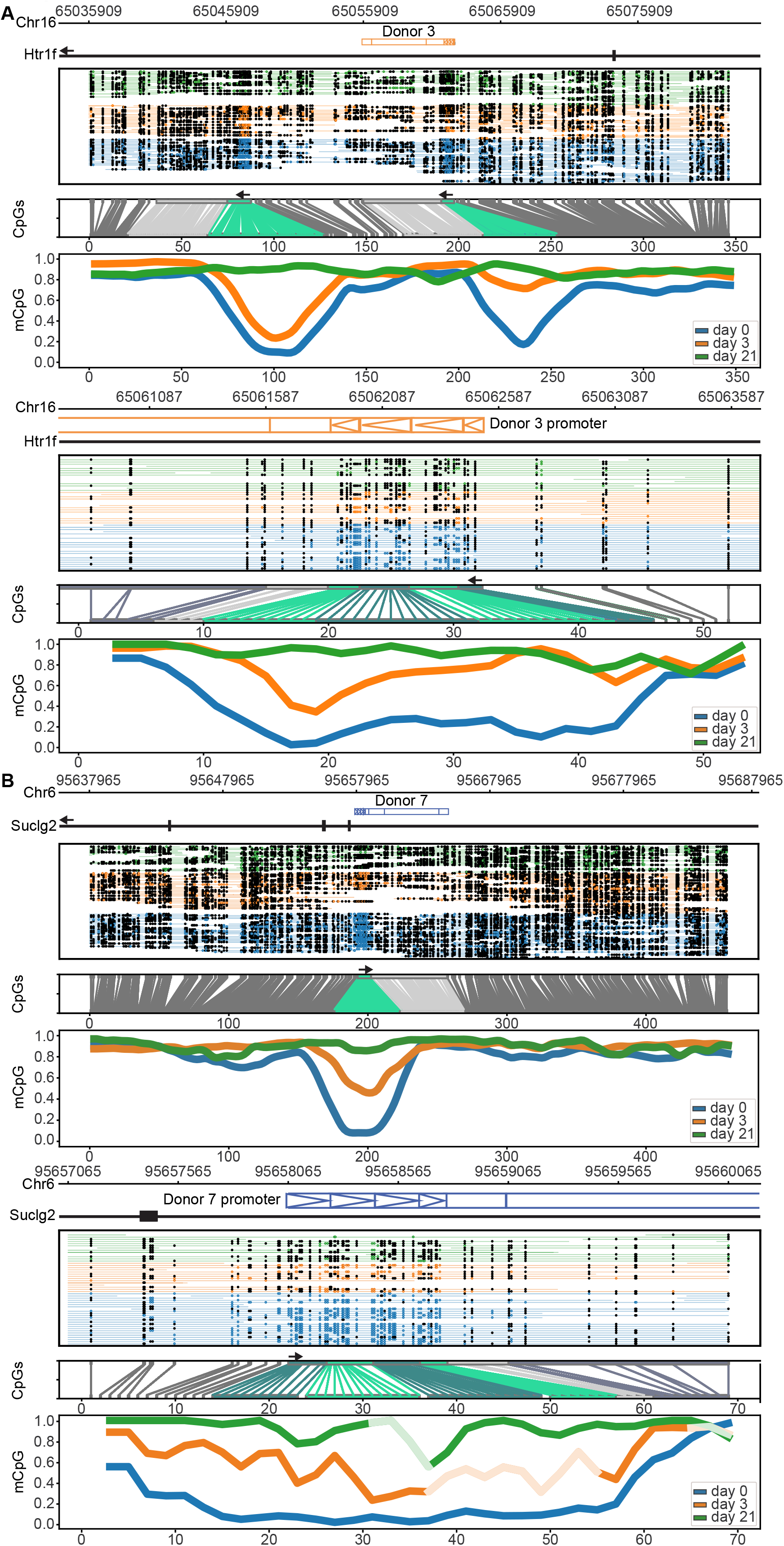
Donor 3 and Donor 7 ONT methylation profiles during mESC differentiation. (*A*) *Top*: Methylation of Donor 3 and surrounding locus. From *top* to *bottom* this figure shows i) the genomic position of Donor 3 in an intron of the *Htr1f* gene on chromosome 16, including 20 kbp up- and downstream of Donor 3, ii) a diagram showing methylated (filled black circles) and unmethylated (unfilled colored circles) CpGs and read (colored lines) coverage per sample, iii) a diagram displaying the correspondence between genome space and CpG space, CpGs belonging to full-length L1 are annotated in light green (promoters) and light grey (ORFs and 3ʹ UTR), iv) the fraction of methylated CpGs for three differentiation time points (d0, d3, d21) in CpG space. *Bottom*: as for upper panels but for promoter of Donor 3, including 1 kbp up- and downstream of the promoter with less smoothing. Monomers in iii) are annotated in light and dark green, unique region in light grey, the start of ORF1 in dark grey/blue. Graphs are shown via a sliding window plot. (*B*) As for (*A*) but for Donor 7 and surrounding locus. The Donor 7 promoter (*bottom* panel (*B*)) smoothed plot lines are colored to appear faded for a short lower confidence region (<20 methylated/demethylated calls within a 30 CpG window).

## DISCUSSION

This study evaluates the mutational capacity of new heritable L1 copies, in terms of their inherent retrotransposition potential and their epigenetic status during development and in adult tissues. We find that loss of 5ʹ monomers by daughter elements relative to their donors consistently diminishes daughter element retrotransposition efficiency. While for individual examples we cannot determine whether this promoter shortening arose due to 5ʹ truncation of the L1 cDNA during retrotransposition (Ostertag et al. 2001; Symer et al. 2002; Zingler et al. 2005), or due to the use of a transcriptional start site internal to the donor L1 promoter, analysis of putative transcription initiator dinucleotides supports the latter hypothesis (Fig. 1C). Moreover, ONT sequencing and methylation analysis revealed that within individual L1 promoters, the 5ʹ-most and 3ʹ-most monomers are more methylated than the internal monomers. Notably, this “smile” pattern of methylation across the L1 promoter is most pronounced in differentiating cells, where repressive mechanisms that compensate for L1 demethylation in pluripotent cells may no longer be present (Walter et al. 2016; Berrens et al. 2017; MacLennan et al. 2017). We propose a model wherein encroaching methylation precludes the use of genome-proximal L1 monomers for initiation of transcription. For donor elements with numerous monomer units which lead to an overall stronger promoter (DeBerardinis and Kazazian 1999), this situation would result in transcription initiation within an internal monomer, generating a shortened mRNA transcript and ultimately giving rise to a daughter insertion with weaker promoter activity and reduced intrinsic retrotransposition potential relative to its donor (Fig. 6). Donor elements with fewer monomers may be entirely transcriptionally silenced under the same conditions (Fig. 4G). The consistent reduction we observe in monomer content between donor and daughter elements may reflect such a phenomenon, although milder instances of promoter shortening may simply indicate use of the first available TSS. Future studies are required to elucidate the direct relationship between the number and position of methylated CpGs within mouse L1 promoters and the ability to initiate transcription.

**Figure 6.**
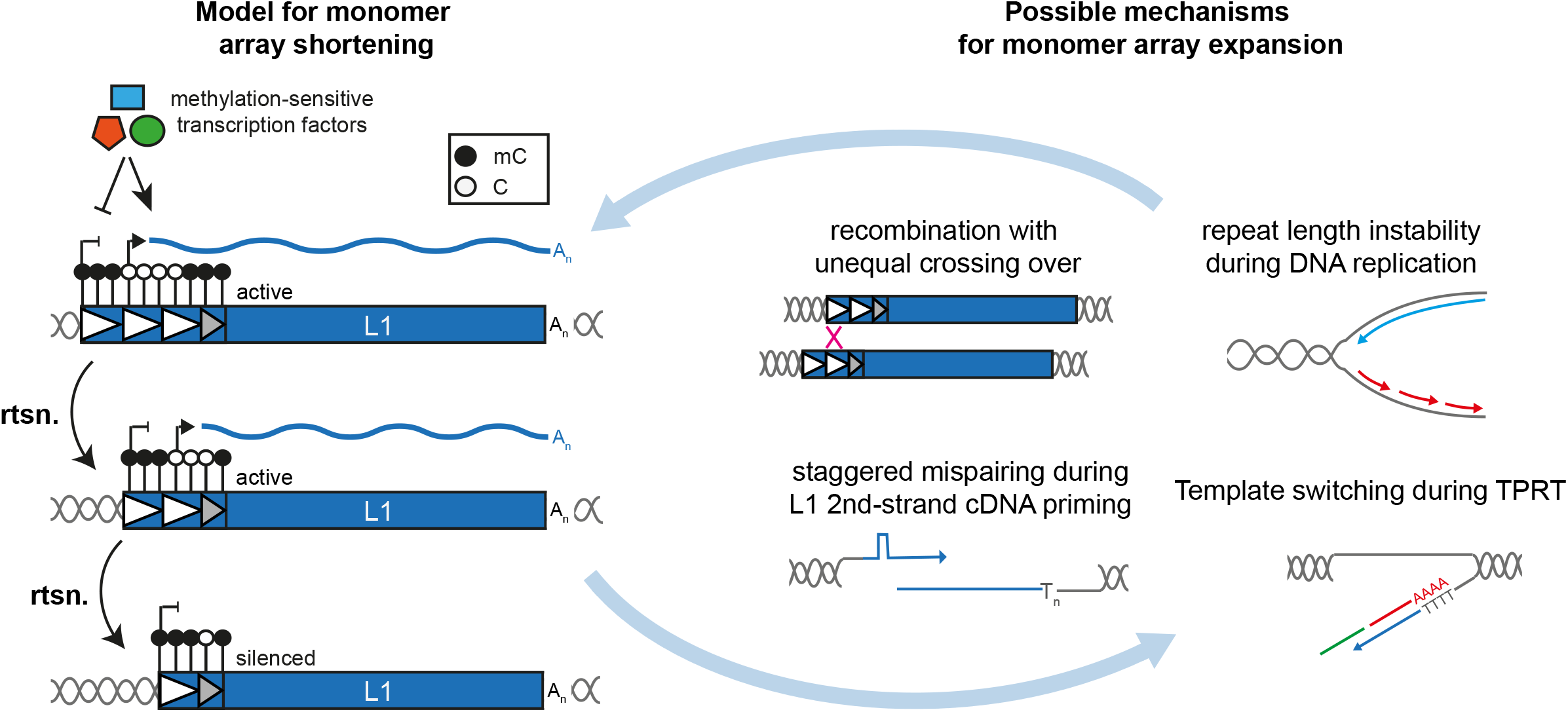
Model for methylation-influenced L1 promoter dynamics during mouse genome evolution. *Left:* Model for mouse L1 monomer array shortening. A lineage of L1 insertions initiating from a donor L1 element harboring three complete promoter monomer units (white triangles) and a 3ʹ partial monomer lacking promoter activity (grey triangle). The use of transcription start sites within internal promoter monomer units (white triangles) is enforced by DNA cytosine methylation (black lollipops) of the 5ʹ- and 3ʹ-most monomers, and resultant inaccessibility to methylation-sensitive transcription factors (colored shapes). With each successive retrotransposition event, loss of 5ʹ monomer sequence reduces the promoter strength and thus the inherent retrotransposition capacity of the daughter element. *Right:* Possible mechanisms for mouse L1 monomer array expansion. Monomer array expansion could occur on extant L1 copies via unequal crossing over between sister chromatids, or due to errors in DNA replication through repeat arrays. Monomers could also be regained concurrent with the generation of a new L1 insertion, either through staggered mispairing of monomer repeats during L1 second-strand cDNA priming, or through either inter- or intra-molecular template switching by the L1 reverse transcriptase.

The repetitive structure and redundant transcription factor binding sites of the mouse L1 promoter may also reduce the impact of mutations which ablate transcription factor binding sites. For example, the impact of a mutation in one YY1 binding site of Insertion 2 may be mitigated due to the intact YY1 binding site located nearby (Fig. 1G). In human L1s, by comparison, loss of the sole YY1 binding site abrogates the capacity to accurately initiate transcription from the +1 position, and thus to ensure a full-length transcript and autonomously mobile daughter insertion (Athanikar et al. 2004). We previously demonstrated the importance of an intact YY1 binding site for efficient methylation of human L1s in pluripotent and differentiated cells (Sanchez-Luque et al. 2019). Mutation of this site leads to hypomethylation of young L1 subfamily members, even in somatic tissues, allowing individual L1 loci to escape epigenetic silencing (Sanchez-Luque et al. 2019). Whether the YY1 binding site plays a similar role in directing methylation of mouse L1s remains to be determined, but it is tempting to speculate that the redundant YY1 binding sites in mouse L1 promoters could enhance the efficiency and coordination of mouse L1 methylation genome-wide and reduce the likelihood of individual loci evading methylation.

In order for L1 lineages to propagate continuously in the mouse genome, diminution of the L1 promoter over successive rounds of retrotransposition must be countered by one or more mechanisms to recover monomer units (Saxton and Martin 1998; Sookdeo et al. 2013; Zhou and Smith 2019). To speculate, recombination with unequal crossing over between sister chromatids could facilitate promoter expansion (Hutchison et al. 1989). Monomer array expansion of extant L1 promoters could also occur during DNA replication, perhaps via DNA slippage or template switching at stalled replication forks (Khristich and Mirkin 2020). Mechanisms to expand monomer arrays concurrent with *de novo* L1 integration include staggered mispairing during priming of second-strand L1 cDNA synthesis (Schichman et al. 1993). Template switching by the L1 reverse transcriptase during TPRT, either between different sites on the same L1 RNA molecule or between two separate L1 RNA molecules, could also account for the duplication of monomer sequences (Gilbert et al. 2005). Indeed, inter-molecular template switching may explain the presence of mouse and rat L1s with “mosaic” 5ʹ UTRs comprised of monomer units from different subfamilies (Adey et al. 1991; Cabot et al. 1997; Hayward et al. 1997). Template switching to non-L1 RNA molecules could also facilitate the capture of novel promoter sequences, which is a common feature in the evolution of rodent L1s as well as chicken CR1 elements (Adey et al. 1991; Hayward et al. 1997; Haas et al. 2001). Overall, we posit that in addition to increasing intrinsic promoter activity, expansion of mouse L1 5ʹ UTR monomer arrays helps maintain retrotransposition competence throughout successive rounds of retrotransposition and across generations, by insulating against monomer loss (Mottez et al. 1986; Loeb et al. 1986) due to potentially methylation-enforced internal transcription initiation as well as 5ʹ truncation and mutation of TF binding sites.

Our precise sequence analysis of endogenous mouse L1 insertions suggests a high fidelity for mouse L1 reverse transcriptase *in vivo*. Comparing the internal sequences of five recent donor/daughter L1 pairs revealed only a single nucleotide misincorporation in 33,374 reverse transcribed bases. Notably, analysis of engineered human L1 insertions generated in HeLa cells revealed 15 misincorporation events in 98,758 reverse transcribed bases, or one error per ∼7,000 bp (Gilbert et al. 2005). Thorough sequence analyses of *de novo* mouse and human L1 insertions generated under the same experimental conditions are required to compare directly the intrinsic fidelity of their reverse transcriptase activities. It is intriguing to speculate that the accuracy of reverse transcription *in vivo* may influence the relative evolutionary success of L1 families in mammalian genomes.

Our locus-specific bisulfite sequencing interrogation of donor and daughter L1 methylation in multiple generations of adult tissues, and in differentiating mESCs *in vitro*, revealed that developmentally active donor L1s and the resultant daughter insertions are generally remethylated in concert with L1 elements genome-wide. *De novo* daughter L1s were methylated even in the somatic and germ tissues of the mosaic animals in which they arose (Fig. 2), likely due to abundant expression of *de novo* DNA methyltransferases in pluripotent cells and early post-implantation embryos (Okano et al. 1998; Chen et al. 2003). This result broadly agrees with a previous study analyzing the methylation status of transgene-derived engineered L1 insertions in cultured cells and transgenic animals (Kannan et al. 2017). It should be noted, however, that Kannan et al. queried the methylation status of heterologous promoters or GFP reporter cassette sequences internal to engineered insertions, rather than the mouse L1 5ʹ UTR. Our ONT analysis across full-length endogenous L1s during differentiation (Fig. 4) demonstrated that the L1 body remains methylated even in pluripotent cells, with the 5ʹ UTR undergoing dynamic methylation changes likely to affect the expression of full-length L1 transcripts and therefore *de novo* retrotransposition events. In addition, the “smile” pattern in methylation across the mouse L1 5ʹ UTR could also lead to an overestimation of mouse L1 promoter methylation, and may impede approaches that analyze only the 5ʹ-most monomers from identifying VM-L1s. Indeed, polyL1Tf_4 is less methylated (<80%) in liver compared to other tissues (>80%) within the same animal, and this methylation pattern is re-established in the F1 but not F2 animal (Fig. 2G). Therefore, it is possible that polyL1Tf_4 is a tissue-specific VM-L1 (Tubio et al. 2014; Schauer et al. 2018; Sanchez-Luque et al. 2019).

In sum, we conclude that the majority of the insertions analyzed here likely arose as a consequence of opportunity provided by genome-wide epigenetic reprogramming during embryonic development (Seisenberger et al. 2012; Hajkova et al. 2002; Cantone and Fisher 2013; Saitou et al. 2012; Abe et al. 2011; Tachibana et al. 2007; Seki et al. 2005). Some exceptional L1s do however escape methylation in a proportion of fully differentiated cells (Fig. S5) or, as for Donor 7, appear methylated in somatic tissues but unmethylated in a large fraction of adult germ cells, while its corresponding daughter insertion and L1 T_F_ monomers genome-wide were highly methylated in the same cells (Fig. 2E,H; Supplemental Fig. S2A,D). Future studies are likely to reveal additional tissue and developmental stage-specific mouse L1 “escapee” loci, with the capacity to evade genome-wide methylation by virtue of their sequence content or surrounding genomic environment.

## METHODS

### Mice

All mouse work was carried out in compliance with the guidelines set forth by the University of Queensland Animal Ethics Committee. Tissues from animals generated for Richardson et al. 2017 were used for this study.

### Cell culture

HeLa-JVM cells were maintained at 37°C and 5% CO_2_ in Dulbecco’s Modified Eagle’s Media (DMEM) (Life Technologies) supplemented with 10% heat-inactivated fetal bovine serum (FBS) (Life Technologies), 1% L-glutamine (Life Technologies) and 1% penicillin-streptomycin (Life Technologies). The cells were passaged every 3-4 days after they achieved confluency of 80-90% using Trypsin 0.25% EDTA (Life Technologies).

E14Tg2a mESCs (ATCC CRL-1821) were cultured on gelatinized tissue culture plates. Plates were coated with 0.1% gelatin (Merck, #ES-006-B) and incubated at 37°C and 5% CO_2_ for 30 min before plating. Cells were passaged at 70-80% confluence every 2-3 days using Trypsin 0.25% EDTA (Life Technologies). mESCs were plated at a density of 1 x 10^5^ cells/ml. Medium was changed every day for optimal growth. mESCs for the retrotransposition assays were cultured in 2i+serum conditions in 1:1 DMEM/F12 media:neurobasal media (Life Technologies) supplemented with 0.5 % N2 and 0.5 % B27 (Life Technologies), 10% FBS (Life Technologies, batch tested), 1% L-glutamine (Life Technologies), 1% penicillin-streptomycin (Life Technologies), 0.1 mM β-mercaptoethanol (Sigma, #M3148), 1 μM PD0325901 (Sigma, #PZ0162) and 3 μM CHIR99021 (Sigma, #SML1046). mESCs for methylation analysis were cultured in 2i+serum conditions as above, 2i+LIF conditions in 1:1 DMEM/F12 media:neurobasal media (Life Technologies) supplemented with N2 and B27 (Life Technologies), 1% L-glutamine (Life Technologies), 1% penicillin-streptomycin (Life Technologies), 0.1% β-mercaptoethanol (Sigma, #M3148), 1 μM PD0325901 (Sigma, #PZ0162), 3 μM CHIR99021 (Sigma, #SML1046) and 1000U/ml ESGRO mLIF (Merck, #ESG1107) and serum+LIF conditions in Knockout DMEM supplemented with 10% FBS (Life Technologies, batch tested), 1% non-essential amino acids (NEAA) (Life Technologies), 1% L-glutamine (Life Technologies), 1% penicillin-streptomycin (Life Technologies), 0.1 mM β-mercaptoethanol (Sigma, #M3148) and 1000U/ml ESGRO mLIF (Merck, #ESG1107).

### Identification of L1 donor/daughter pairs

We previously identified 11 *de novo* endogenous L1 T_F_ insertions among 85 C57BL6/J mice belonging to multi-generation pedigrees, as well as 6 unfixed polymorphic L1 T_F_ insertions absent from the C57BL6/J reference genome and differentially present/absent among these pedigrees (Richardson et al. 2017). We traced the majority of heritable insertions to pluripotent embryonic cells, evidenced by shared somatic/germline mosaicism of the founder mouse, and early primordial germ cells (PGCs), evidenced by germline-restricted mosaicism across both testes of male founder mice. Analysis of unique L1 3ʹ transduced sequences (Xing et al. 2006; Goodier et al. 2000; Pickeral et al. 2000; Moran et al. 1999; Holmes et al. 1994) allowed us to identify the source (donor) L1 loci responsible for three offspring (daughter) *de novo* insertions (one early embryonic, two early PGC) and two unfixed polymorphic daughter insertions.

### DNA extraction

Genomic DNA from mouse tissue was extracted as previously described (Richardson et al. 2017). Genomic DNA from cultured cells was extracted using DNeasy Blood & Tissue kit (Qiagen) according to the manufacturer’s protocol. The DNA concentration was determined by Qubit 3.0 Fluorometer (Invitrogen) using the Qubit dsDNA HS Assay Kit (Invitrogen) following the manufacturer’s instructions.

High molecular weight (HMW) genomic DNA from cultured cells was extracted using the Nanobind CBB Big DNA Kit (Circulomics) following the manufacturer’s instructions.

### Generation of mouse L1 reporter constructs

DNA sequences corresponding to the different donor/daughter mouse L1 elements were amplified from genomic DNA using Expand Long Range dNTPack (Roche). Reaction mixes contained 5 μl 5x Expand Long Range Buffer with 12.5 mM MgCl_2_, 1.25 μl dNTP Mix (dATP, dCTP, dGTP, dTTP at 10 mM each), 1.25 μl DMSO (100%), 1 μl primer mix (50 μM of each primer), 0.35 μl Expand Long Range Enzyme Mix (5 U/μl), 4-10 ng genomic DNA template and molecular grade water up to a total volume of 25 μl. PCRs were performed with the following cycling conditions: 92°C for 3 min, 10 cycles of 92°C for 30 sec, 56-60°C for 30 sec, and 68°C for 5-7.5 min; 25 cycles of 92°C for 30 sec, 56-60°C for 30 sec, and 68°C for 5-7 min plus 20 sec/cycle elongation for each successive cycle, followed by 68°C for 10 min. Primers (Supplemental Table 1) introduced a *NotI* restriction site at the L1 5ʹ end. Full-length L1 elements were then cloned into pGEMT Easy Vector (Promega) according to the manufacturer’s instructions. Ligations were incubated overnight at 4°C. Ligation reactions were transformed using One Shot TOP10 chemically competent *E. coli* (Invitrogen) according to the manufacturer’s instructions. Blue/white screening was performed using LB/ampicillin/IPTG/X-Gal plates. 3-5 positive colonies per L1 element were chosen for Miniprep culture and plasmid DNA was isolated using QIAprep Spin Miniprep Kit (Qiagen) according to the manufacturer’s instructions. At least three clones per element were capillary sequenced using L1 sequencing primers (Supplemental Table 1). Sequences from at least three clones of the same L1 element were compared to each other to identify PCR-induced mutations. L1 elements were then reconstructed by combination of non-mutated fragments from different clones using restriction enzymes (New England Biolabs) cutting within the L1 sequence.

The L1 3ʹend fragment was produced by PCR amplification from plasmid DNA using a forward primer upstream of a *HindIII*-site in ORF2 and a reverse primer introducing an *SbfI*-site at the end of the L1 sequence thereby removing the polyadenylation signal (AATAAA) and the G-rich region (GRR). The GRR was amplified using primers that introduce an *AgeI*- and *PacI*-site on each site of the GRR and removing the polyadenylation signal. All restriction enzymes used for cloning were obtained from New England Biolabs. Restriction digests were performed according to the manufacturer’s instructions (New England Biolabs). Reactions were purified using agarose gel electrophoresis and target fragments were excised and purified using QIAquick and MinElute Gel Extraction Kits (Qiagen) according to the manufacturer’s protocol.

A modified version of the previously described pTN201 construct (Naas et al. 1998) in which the L1 3ʹ UTR GRR is located downstream, rather than upstream, of the NEO indicator cassette was used as a backbone to generate L1 reporter constructs (Richardson et al., in preparation). L1spa was removed from the pCEP4 backbone using *NotI* and *SbfI*. The pCEP4 backbone was dephosphorylated using Calf Intestinal Alkaline Phosphatase (CIP) (New England Biolabs) according to the manufacturer’s instructions. The backbone and multiple fragments from the target L1 (Insertion 2, Insertion 5, Insertion 7, polyL1Tf_3, polyL1Tf_4, Donor 2, Donor 5, Donor 7, Donor 3, Donor 4) were combined in a single ligation reaction using T4 DNA Ligase (New England Biolabs) according to the manufacturer’s instructions and incubated overnight at 16°C. Ligations were transformed using One Shot TOP10 chemically competent *E. coli* (Invitrogen) according to the manufacturer’s instructions. Plasmid DNA of positive clones was obtained using QIAprep Spin Miniprep Kit (Qiagen). The absence of mutations was verified by capillary sequencing. The GRR of L1spa was replaced by the respective GRR of each L1 element using *AgeI* and *PacI*. The backbone and GRR were ligated and transformed as described above. Plasmid DNA for the retrotransposition assays was obtained using Plasmid Maxi kit (Qiagen). The absence of mutations was verified by capillary sequencing. Each construct was built with and without a CMV promoter upstream of the L1 element.

Insertion 2 has a C to T mutation in the YY1 binding motif in its first monomer upstream of the unique region. To fix the mutation, a reverse primer (Insertion_2_YY1_fix_R) overlapping the mutation with the correct YY1 binding motif sequence was designed (Supplemental Table 1). The promoter was then amplified from the construct prepared for the retrotransposition assay using Q5 High-Fidelity 2x Master Mix (New England Biolabs). Primers and annealing temperatures are listed in Supplemental Table 1. Reaction mixes contained 12.5 μl Q5 High-Fidelity 2x Master Mix, 1.25 μl primer mix (10 μM of each primer), 2 ng plasmid DNA template and molecular grade water up to a total volume of 25 μl. PCRs were performed using the following conditions: 98°C for 2 min, 35 cycles of 98°C for 10 sec, 62-72°C for 30 sec, and 72°C for 1 min, followed by 72°C for 2 min. Insertion_2_FL_F was used as a forward primer. The PCR fragment was then digested with *NotI* and *XmaI*. pCEP4ΔCMV-mneoI-G4-Ins2 was used as a backbone and digested with *NotI* and *SbfI*. The backbone was dephosphorylated using Calf Intestinal Alkaline Phosphatase (CIP) (New England Biolabs) according to the manufacturer’s instructions. The rest of the Insertion 2 element was obtained by digesting pCEP4ΔCMV-mneoI-G4-Ins2 with *XmaI* and *SbfI*. The three fragments were then ligated and transformed as described above. Plasmid DNA of positive clones was obtained using Plasmid Maxi Kit (Qiagen). The absence of mutations was verified by capillary sequencing.

### mESC differentiation

E14Tg2a mESCs (ATCC CRL-1821) were differentiated into embryoid bodies (EBs) as previously described (Behringer et al. 2016). Hanging drop culture was used to generate EBs of comparable dimension. Bacteriological-grade 10 cm plates were prepared by adding 15 ml 1x PBS (Invitrogen) to each plate and incubating them at 37°C and 5% CO_2_. mESCs cultured in serum+LIF conditions were harvested at 60-70% confluency by adding Trypsin 0.25% EDTA (Life Technologies). The trypsinized cells were resuspended in EB medium (Knockout DMEM (Life Technologies, 10% FBS (Life Technologies, batch tested), 1% non-essential amino acids (NEAA) (Life Technologies), 1% L-glutamine (Life Technologies), 1% penicillin-streptomycin (Life Technologies), 0.1% β-mercaptoethanol (Sigma, #M3148)) and centrifuged at 400 xg for 5 mins. Supernatant was removed and cells were resuspended in EB medium. The cell concentration was determined using the TC20 Automated Cell Counter (Bio-Rad) and the cell suspension was diluted to 3 x 10^4^ cells/ml. 30 μl cell suspension drops (1000 cells/drop) were distributed inside the lid of the previously prepared bacteriological-grade 10 cm plates. The lid was carefully placed back on the base containing 1x PBS (Invitrogen). The EBs were incubated for three days. On day three, EBs were collected from lids and transferred to six-well ultra-low adherence plates (Corning). 2 ml EB media were added to each well and the EBs were incubated for another three days. Six day old EBs were plated onto six-well plates coated with 0.1% gelatin (Merck) in EB medium. After 24 h, the EB medium was replaced by differentiation medium (Knockout DMEM (Life Technologies), 1% non-essential amino acids (NEAA) (Life Technologies), 1% L-glutamine (Life Technologies), 1% penicillin-streptomycin (Life Technologies), 0.1% β-mercaptoethanol (Sigma, #M3148)). The cells were differentiated for two weeks. Medium was changed every other day and cells for DNA extraction were collected every three days.

### Immunostaining

Immunostaining was performed on day 7 (24 h after plating of EBs) and day 21 of mESC differentiation. EBs were plated on gelatinized coverslips in twelve-well plates. Coverslips harboring cells were rinsed with 1x PBS (Invitrogen) and fixed in 4% paraformaldehyde in PBS for 20 mins at room temperature. Coverslips were washed twice with 1x PBS for 5 min followed by 1 min incubation in permeabilisation buffer PBT (1x PBS, 0.5% Triton x100 (Sigma)). Coverslips were washed in 1x PBS for 10 min and blocked in PBT containing 10% normal donkey serum (blocking buffer) for 1 h at room temperature. Primary antibodies (β-tubulin, Rabbit IgG (Sigma, #T2200), 1:500; Α-fetoprotein (AFP), Goat IgG (R&D Systems, #AF5369), 1:200; Smooth muscle actin, Mouse IgG (Life Technologies, #14976080), 1:500) were diluted in blocking buffer and incubated for 1 h at room temperature, then washed with 1x PBS. Secondary antibodies (Alexa Fluor 647 Donkey Anti-Rabbit IgG (Jackson ImmunoResearch, #711-606-152), 1:500; Cy3 Donkey Anti-Goat IgG (Jackson ImmunoResearch, #715-165-150), 1:200; Alexa Fluor 488 Donkey Anti-Mouse IgG (Jackson ImmunoResearch, #715-546-150), 1:500) were diluted in blocking buffer and incubated for 1 h at room temperature, then washed with 1x PBS. Cells were stained with H33258 (1:1000 in 1x PBS) for 5 min and washed with 1x PBS for 10 min. PermaFluor mounting media (Thermo Fisher Scientific, #TA-030-FM) was added to glass chamber slides and coverslips were carefully placed to avoid the creation of bubbles and dried overnight at room temperature. Cells were imaged on a spinning-disk confocal system (Marianas; 3I, Inc.) consisting of an Axio Observer Z1 (Carl Zeiss) equipped with a CSU-W1 spinning-disk head (Yokogawa Corporation of America), ORCA-Flash4.0 v2 sCMOS camera (Hamamatsu Photonics) and a 20x 0.8 NA PlanApo objective. Image acquisition was performed using SlideBook 6.0 (3I, Inc). Image processing and analysis was done using Image J 1.52 software.

### Retrotransposition assay

Retrotransposition assays in HeLa-JVM cells were performed as previously described (Kopera et al. 2016) with some minor modifications. HeLa-JVM cells were grown in DMEM complete medium. To assay L1 retrotransposition of L1 donor/daughter pairs, HeLa-JVM were seeded at a density of 5 x 10^3^ cells/well in six-well tissue culture plates. To assay Insertion 2-YY1-fixed, HeLa-JVM were seeded at a density of 1 x 10^4^ cells/well. 14-16 h after plating, cells were transfected with L1 reporter constructs using 4 μl FuGENE HD transfection reagent (Promega), 96 μl Opti-MEM (Life Technologies) and 1 μg plasmid DNA per well. Transfection efficiency was determined in parallel by preparing transfection mixes containing 4 μl FuGENE HD transfection reagent (Promega), 96 μl Opti-MEM (Life Technologies), 0.5 μg L1 expression plasmid and 0.5 μg pCEP4-eGFP. 100 μl transfection mixture was added to each well containing 2 mL DMEM complete medium. The plates were incubated at 37°C and 5% CO_2_. The transfection was stopped by replacing the medium 24 h post-transfection. Transfection efficiency was determined 72 h post-transfection. pCEP4-eGFP co-transfected wells were trypsinized and cells were collected from each well and centrifuged at 400 xg for 5 min. Cell pellets were resuspended in 300-500 μl 1x PBS (Invitrogen). 10 μl Propidiumiodide (Life Technologies) were added to cell suspensions. The number of eGFP-positive cells was determined using a CytoFLEX flow cytometer (Beckman Coulter). The percentage of eGFP-positive cells was used to normalize the G418-resistant colony counts for each L1 reporter construct (Kopera et al. 2016). Geneticin/G418 (400 μg/ml) (Thermo Fisher Scientific) selection was started 3 days post-transfection and performed for 12 days. G418-resistant foci were washed with 1x PBS and fixed using 2% Formaldehyde/0.2% Glutaraldehyde in 1x PBS (Sigma-Aldrich) fixing solution at room temperature for 30 min. Staining was done using 0.1% Crystal Violet solution (Sigma-Aldrich) at room temperature for 10 min. Foci were counted in each well. Three biological replicates, each containing three technical replicates per L1 construct were done.

L1 retrotransposition assays in E14Tg2a mESCs (ATCC CRL-1821) were performed as previously described (MacLennan et al. 2017) by plating 4 x 10^5^ cells/well in gelatin-coated six-well plates in antibiotic-free 2i+serum conditions. After 18 h, the cells were transfected using Lipofectamine 2000 (Invitrogen) according to the manufacturer’s instructions. Briefly, 1 μg of plasmid DNA was diluted in 125 μl Opti-MEM (Life Technologies). 4 μl Lipofectamine 2000 were diluted in 125 μl Opti-MEM (Life Technologies) and incubated for 5 min at room temperature. The DNA/Opti-MEM and Lipofectamine/Opti-MEM mixtures were combined and incubated for 20 min at room temperature. 250 μl transfection mixture was added to each well. The medium was replaced after 8 h. Transfection efficiency was determined in parallel by preparing transfection mixes containing 0.5 μg L1 expression plasmid and 0.5 μg pCEP4-eGFP. Transfection efficiency was determined 24 h after transfection as described for HeLa-JVM cells. 24 h after transfection, the mESCs were passaged into gelatin-coated 10 cm dishes using Trypsin 0.25% EDTA (Life Technologies). G418 (Invitrogen) selection (200 μg/ml) was started 24 h after passaging and continued for 12 days. Drug-resistant colonies were fixed, stained and counted as described for HeLa-JVM cells. Three biological replicates, each containing two technical replicates per L1 construct were done.

### Locus-specific bisulfite sequencing

Bisulfite conversion was performed with 200-400 ng input genomic DNA using the EZ DNA Methylation-Lightning Kit (Zymo Research), following the manufacturer’s instructions. DNA was eluted in 10 μl Elution Buffer. Primers used for target amplification are listed in Supplemental Table 1.

PCRs were performed using MyTaq HS DNA Polymerase (Bioline). Reaction mixes contained 5 μl 5x MyTaq Reaction Buffer, 0.5 μl primer mix (25 μM of each primer), 0.2 μl MyTaq HS DNA Polymerase, DMSO at a final concentration of 0.1%, 2 μl bisulfite converted DNA template and molecular grade water up to a total volume of 25 μl. PCRs were performed using the following conditions: 95°C for 2 min, 40 cycles of 95°C for 30 sec, 54°C for 30 sec, and 72°C for 30 sec, followed by 72°C for 5 min. PCR products were run on a 2% agarose gel, excised and purified using the MinElute Gel Extraction Kit (Qiagen) according to the manufacturer’s instructions. Illumina libraries were constructed using the NEBNext Ultra II DNA Library Prep Kit (New England Biolabs) following the manufacturer’s instructions. Library quantity was determined using the Bioanalyzer DNA 1000 chip (Agilent Technologies) according to the manufacturer’s instructions. Barcoded libraries were pooled equimolar and sequenced on an Illumina MiSeq platform using a MiSeq Reagent Kit v3 (600-cycle). 50% PhiX Control v3 (Illumina) was used as spike-in and cluster density was intended to be ∼800-1000K/mm2.

Analysis was performed as previously described (Schauer et al. 2018). Paired-end sequencing reads (>200 bp) were assembled into contigs using FLASH (Magoč and Salzberg 2011) with default parameters. Contigs per sample were identified by primer sequences carried at their termini. Unconverted amplicon sequences are listed in Supplemental Table 1.

Per sample, 50 reads were randomly selected and analyzed using QUMA (QUantification tool for Methylation Analysis) (Kumaki et al. 2008) with default parameters, plus requiring strict CpG recognition and, in the case of L1 methylation, excluding identical bisulfite sequences.

### Nanopore sequencing

Purity of HMW DNA was determined using a NanoDrop One Spectrophotometer (Thermo Fisher Scientific) according to the manufacturer’s instructions. The DNA concentration was determined by Qubit 3.0 Fluorometer (Invitrogen) using the Qubit dsDNA HS Assay Kit (Invitrogen) and on a TapeStation System (Agilent Technologies). ONT sequencing libraries were created using 1D Ligation (SQK-LSK109), sheared to create an average fragment size of ∼10 kb and sequenced at the Kinghorn Centre for Clinical Genomics at the Garvan Institute of Medical Research (Darlinghurst, NSW, Australia) on an ONT PromethION platform.

Bases were called using Guppy version 3.2.10 (Oxford Nanopore Technologies) and aligned to the reference genome build mm10 using minimap2 version 2.17 (Li 2018) and samtools version 1.3 (Li et al. 2009). Reads were indexed and per-CpG methylation calls generated using nanopolish version 0.13.2 (Simpson et al. 2017). Methylation likelihood data were sorted by position and indexed using tabix version 1.10.2 (Li 2011).

Reference L1 locations were derived from the RepeatMasker (http://www.repeatmasker.org/).out track files available for mm10 from the UCSC Genome Browser (Kent 2002). As full-length mouse L1s are often broken into multiple adjacent annotations when present on the - strand of the genome, we merged adjacent similarly oriented L1s prior to analysis. L1s were considered if annotated as greater than 6000 bp in length. Monomers were counted using a python script which considers the best alignment between a library of known monomer sequences and a target transposable element using exonerate (Slater and Birney 2005), records and masks the monomer alignment, and repeats the process until no further monomer alignments are present. Methylation results were annotated per-subfamily and per-monomer count and plotted using these categories. Methylation statistics for reference L1s were generated using the “segmeth” function in methylartist (https://github.com/adamewing/methylartist) plotted using the “segplot” function. Reads mapping completely within L1s were excluded from the reference L1 methylation analysis via the “--exclude_ambiguous” option in methylartist segmeth to negate the contribution of ambiguous mappings. Plots categorized by monomer count (Fig. 4D) were limited to reads spanning the entire segment with the addition of the “--spanning_only” argument to segmeth. Methylation plots for individual L1s and differentially methylated regions (DMRs) (Fig. 5) were created using the methylartist “locus” function. Composite per-element methylation plots (Fig. 4F-G; Supplemental Fig. S5,S7) were created using the methylartist “composite” function.

## DATA ACCESS

ONT DNA sequencing raw data has been deposited in the NCBI BioProject database under the Project ID PRJNA763783.

## ACKNOWLEDGEMENTS

The authors thank Dr. John V. Moran, Dr. Ian R. Adams and members of the Faulkner lab for helpful advice and discussion. This research was supported by an Australian Government Research Training Program Scholarship and a Mater Research Frank Clair Scholarship awarded to P.G., a University of Queensland Research Training Program Scholarship and Commonwealth Scientific and Industrial Research Organisation Postgraduate Top-Up Scholarship awarded to M.L., the People Programme (Marie Curie Actions) of the European Union Seventh Framework Program (FP7/2007-2013) under REA grant agreement PIOF-GA-2013-623324 awarded to F.J.S.-L. and the Australian National Health and Medical Research Council (NHMRC) and Australian Research Council (ARC) Dementia Research Development Fellowship GNT1108258 awarded to G.O.B.. This study was funded by the Australian NHMRC (GNT1125645, GNT1138795 and GNT1173711 to G.J.F.; GNT1173476 to S.R.R.), The ARC (DP200102919 to G.J.F. and S.R.R.), a CSL Centenary Fellowship to G.J.F., an Advance Queensland Women’s Academic Fund Maternity Funding award to S.R.R., the Mater Research Strategic Grant for Outstanding Women to S.R.R, and the Mater Foundation (Equity Trustees / AE Hingeley, QFC Thomas George and KC BM Thomson Trusts). We acknowledge the TRI flow cytometry core for technical assistance and equipment. We acknowledge QBI Advanced Microscopy Facility for technical assistance and equipment, supported by ARC LIEF grant LE130100078.

## AUTHOR CONTRIBUTIONS

P.G., D.C., F.J.S.-L., G.O.B. and S.R.R. designed and performed experiments. P.G., M.L., A.D.E. and G.J.F. performed bioinformatics analyses. P.G. and S.R.R. conceived the study. P.G., G.J.F. and S.R.R. wrote the manuscript. All authors commented on and approved the final manuscript.

## REFERENCES

Abe M, Tsai SY, Jin S-G, Pfeifer GP, Szabó PE. 2011. Sex-Specific Dynamics of Global Chromatin Changes in Fetal Mouse Germ Cells ed. E. Ballestar. PLoS One 6: e23848.

Adey NB, Schichman SA, Graham DK, Peterson SN, Edgell MH, Hutchison CA. 1994. Rodent L1 evolution has been driven by a single dominant lineage that has repeatedly acquired new transcriptional regulatory sequences. Mol Biol Evol 11: 778–89.

Adey NB, Schichman SA, Hutchison CA, Edgell MH. 1991. Composite of A and F-type 5′ terminal sequences defines a subfamily of mouse LINE-1 elements. J Mol Biol 221: 367–373.

Akagi K, Li J, Stephens RM, Volfovsky N, Symer DE. 2008. Extensive variation between inbred mouse strains due to endogenous L1 retrotransposition. Genome Res 18: 869–80.

Athanikar JN, Badge RM, Moran JV. 2004. A YY1-binding site is required for accurate human LINE-1 transcription initiation. Nucleic Acids Res 32: 3846–3855.

Beck CR, Collier P, Macfarlane C, Malig M, Kidd JM, Eichler EE, Badge RM, Moran JV. 2010. LINE-1 Retrotransposition Activity in Human Genomes. Cell 141: 1159–1170.

Behringer R, Gertsenstein M, Nagy KV, Nagy A. 2016. Differentiating mouse embryonic stem cells into embryoid bodies by hanging-drop cultures. Cold Spring Harb Protoc 2016: 1073–1076.

Berrens RV, Andrews S, Spensberger D, Santos F, Dean W, Gould P, Sharif J, Olova N, Chandra T, Koseki H, et al. 2017. An endosiRNA-Based Repression Mechanism Counteracts Transposon Activation during Global DNA Demethylation in Embryonic Stem Cells. Cell Stem Cell 21: 694–703.e7.

Bertozzi TM, Ferguson-Smith AC. 2020. Metastable epialleles and their contribution to epigenetic inheritance in mammals. Semin Cell Dev Biol 97: 93–105.

Besse S, Allamand V, Vilquin J-T, Li Z, Poirier C, Vignier N, Hori H, Guénet J-L, Guicheney P. 2003. Spontaneous muscular dystrophy caused by a retrotransposal insertion in the mouse laminin α2 chain gene. Neuromuscul Disord 13: 216–222.

Bourc’his D, Bestor TH. 2004. Meiotic catastrophe and retrotransposon reactivation in male germ cells lacking Dnmt3L. Nature 431: 96–99.

Cabot EL, Angeletti B, Usdin K, Furano A V. 1997. Rapid evolution of a young L1 (LINE-1) clade in recently speciated rattus taxa. J Mol Evol 45: 412–423.

Cantone I, Fisher AG. 2013. Epigenetic programming and reprogramming during development. Nat Struct Mol Biol 20: 282–9.

Carninci P, Sandelin A, Lenhard B, Katayama S, Shimokawa K, Ponjavic J, Semple CAM, Taylor MS, Engström PG, Frith MC, et al. 2006. Genome-wide analysis of mammalian promoter architecture and evolution. Nat Genet 38: 626–635.

Casavant NC, Hardies SC. 1994. The dynamics of murine LINE-1 subfamily amplification. J Mol Biol 241: 390–397.

Castro-Diaz N, Ecco G, Coluccio A, Kapopoulou A, Yazdanpanah B, Friedli M, Duc J, Jang SM, Turelli P, Trono D. 2014. Evolutionally dynamic L1 regulation in embryonic stem cells. Genes Dev 28: 1397–1409.

Cheetham SW, Kindlova M, Ewing AD. 2021. Methylartist: Tools for Visualising Modified Bases from Nanopore Sequence Data. bioRxiv 2021.07.22.453313.

Chen T, Ueda Y, Dodge JE, Wang Z, Li E. 2003. Establishment and maintenance of genomic methylation patterns in mouse embryonic stem cells by Dnmt3a and Dnmt3b. Mol Cell Biol 23: 5594–605.

Crooks GE, Hon G, Chandonia JM, Brenner SE. 2004. WebLogo: A sequence logo generator. Genome Res 14: 1188–1190.

de la Rica L, Deniz Ö, Cheng KCL, Todd CD, Cruz C, Houseley J, Branco MR. 2016. TET-dependent regulation of retrotransposable elements in mouse embryonic stem cells. Genome Biol 17: 234.

DeBerardinis RJ, Goodier JL, Ostertag EM, Kazazian HH. 1998. Rapid amplification of a retrotransposon subfamily is evolving the mouse genome. Nat Genet 20: 288–290.

DeBerardinis RJ, Kazazian HH. 1999. Analysis of the Promoter from an Expanding Mouse Retrotransposon Subfamily. Genomics 56: 317–323.

Deniz Ö, Frost JM, Branco MR. 2019. Regulation of transposable elements by DNA modifications. Nat Rev Genet 20: 417–431.

Dombroski BA, Mathias SL, Nanthakumar E, Scott AF, Kazazian HH. 1991. Isolation of an active human transposable element. Science 254: 1805–8.

Doucet AJ, Hulme AE, Sahinovic E, Kulpa DA, Moldovan JB, Kopera HC, Athanikar JN, Hasnaoui M, Bucheton A, Moran JV, et al. 2010. Characterization of LINE-1 Ribonucleoprotein Particles ed. G.S. Barsh. PLoS Genet 6: e1001150.

Doucet AJ, Wilusz JE, Miyoshi T, Liu Y, Moran JV. 2015. A 3’ Poly(A) Tract Is Required for LINE-1 Retrotransposition. Mol Cell 60: 728–741.

Elmer JL, Hay AD, Kessler NJ, Bertozzi TM, Ainscough EAC, Ferguson-Smith AC. 2021. Genomic properties of variably methylated retrotransposons in mouse. Mob DNA 12: 1–16.

Ergun S, Buschmann C, Heukeshoven J, Dammann K, Schnieders F, Lauke H, Chalajour F, Kilic N, Stratling WH, Schumann GG. 2004. Cell Type-specific Expression of LINE-1 Open Reading Frames 1 and 2 in Fetal and Adult Human Tissues. J Biol Chem 279: 27753–27763.

Ewing AD, Smits N, Sanchez-Luque FJ, Faivre J, Brennan PM, Richardson SR, Cheetham SW, Faulkner GJ. 2020. Nanopore Sequencing Enables Comprehensive Transposable Element Epigenomic Profiling. Mol Cell 80: 915–928.e5.

Feng Q, Moran JV, Kazazian HH, Boeke JD. 1996. Human L1 retrotransposon encodes a conserved endonuclease required for retrotransposition. Cell 87: 905–916.

Flasch DA, Macia Á, Sánchez L, Ljungman M, Heras SR, García-Pérez JL, Wilson TE, Moran JV. 2019. Genome-wide de novo L1 Retrotransposition Connects Endonuclease Activity with Replication. Cell 177: 837–851.e28.

Furano A V, Robb SM, Robb FT. 1988. The structure of the regulatory region of the rat L1 (L1Rn, long interspersed repeated) DNA family of transposable elements. Nucleic Acids Res 16: 9215–31.

Gagnier L, Belancio VP, Mager DL. 2019. Mouse germ line mutations due to retrotransposon insertions. Mob DNA 10: 15.

Gardner EJ, Lam VK, Harris DN, Chuang NT, Scott EC, Stephen Pittard W, Mills RE, Devine SE. 2017. The mobile element locator tool (MELT): Population-scale mobile element discovery and biology. Genome Res 27: 1916–1929.

Gilbert N, Lutz S, Morrish TA, Moran JV. 2005. Multiple Fates of L1 Retrotransposition Intermediates in Cultured Human Cells. Mol Cell Biol 25: 7780–7795.

Goodier JL. 2016. Restricting retrotransposons: a review. Mob DNA 7: 16.

Goodier JL, Ostertag EM, Du K, Kazazian HH. 2001. A novel active L1 retrotransposon subfamily in the mouse. Genome Res 11: 1677–85.

Goodier JL, Ostertag EM, Kazazian HH. 2000. Transduction of 3’-flanking sequences is common in L1 retrotransposition. Hum Mol Genet 9: 653–7.

Greenberg MVC, Bourc’his D. 2019. The diverse roles of DNA methylation in mammalian development and disease. Nat Rev Mol Cell Biol 20: 590–607.

Grimaldi G, Skowronski J, Singer MF. 1984. Defining the beginning and end of KpnI family segments. EMBO J 3: 1753–9.

Haas NB, Grabowski JM, North J, Moran JV, Kazazian HH, Burch JBE. 2001. Subfamilies of CR1 non-LTR retrotransposons have different 5′UTR sequences but are otherwise conserved. Gene 265: 175– 183.

Hajkova P, Erhardt S, Lane N, Haaf T, El-Maarri O, Reik W, Walter J, Surani MA. 2002. Epigenetic reprogramming in mouse primordial germ cells. Mech Dev 117: 15–23.

Han JS, Boeke JD. 2004. A highly active synthetic mammalian retrotransposon. Nature 429: 314–318.

Hardies SC, Wang L, Zhou L, Zhao Y, Casavant NC, Huang S. 2000. Line-1 (L1) lineages in the mouse. Mol Biol Evol 17: 616–628.

Hata K, Sakaki Y. 1997. Identification of critical CpG sites for repression of L1 transcription by DNA methylation. Gene 189: 227–34.

Hayward BE, Zavanelli M, Furano A V. 1997. Recombination creates novel L1 (LINE-1) elements in Rattus norvegicus. Genetics 146: 641–54.

Hohjoh H, Singer MF. 1996. Cytoplasmic ribonucleoprotein complexes containing human LINE-1 protein and RNA. EMBO J 15: 630–9.

Holmes SE, Dombroski BA, Krebs CM, Boehm CD, Kazazian HH. 1994. A new retrotransposable human L1 element from the LRE2 locus on chromosome 1q produces a chimaeric insertion. Nat Genet 7: 143–148.

Holmes SE, Singer MF, Swergold GD. 1992. Studies on p40, the leucine zipper motif-containing protein encoded by the first open reading frame of an active human LINE-1 transposable element. J Biol Chem 267: 19765–8.

Hutchison IC, Hardies S, Loeb D, Shehee W, Edgell M. 1989. LINEs and related retroposons: long interspersed repeated sequences in the eukayotic genome. In Mobile DNA (eds. D. Berg and M. Howe), pp. 593–617, ASM Press.

Jacobs FMJ, Greenberg D, Nguyen N, Haeussler M, Ewing AD, Katzman S, Paten B, Salama SR, Haussler D. 2014. An evolutionary arms race between KRAB zinc-finger genes ZNF91/93 and SVA/L1 retrotransposons. Nature 516: 242–5.

Kannan M, Li J, Fritz SE, Husarek KE, Sanford JC, Sullivan TL, Tiwary PK, An W, Boeke JD, Symer DE. 2017. Dynamic silencing of somatic L1 retrotransposon insertions reflects the developmental and cellular contexts of their genomic integration. Mob DNA 8: 8.

Kazachenka A, Bertozzi TM, Sjoberg-Herrera MK, Walker N, Gardner J, Gunning R, Pahita E, Adams S, Adams D, Ferguson-Smith AC. 2018. Identification, Characterization, and Heritability of Murine Metastable Epialleles: Implications for Non-genetic Inheritance. Cell 175: 1259–1271.e13.

Kent WJ. 2002. BLAT---The BLAST-Like Alignment Tool. Genome Res 12: 656–664.

Khazina E, Weichenrieder O. 2018. Human LINE-1 retrotransposition requires a metastable coiled coil and a positively charged N-terminus in L1ORF1p. Elife 7: e34960.

Khazina E, Weichenrieder O. 2009. Non-LTR retrotransposons encode noncanonical RRM domains in their first open reading frame. Proc Natl Acad Sci U S A 106: 731–6.

Khristich AN, Mirkin SM. 2020. On the wrong DNA track: Molecular mechanisms of repeat-mediated genome instability. J Biol Chem 295: 4134–4170.

Kim J do, Kim J. 2009. YY1’s longer DNA-binding motifs. Genomics 93: 152–158.

Kingsmore SF, Giros B, Suh D, Bieniarz M, Caron MG, Seldin MF. 1994. Glycine receptor β–subunit gene mutation in spastic mouse associated with LINE–1 element insertion. Nat Genet 7: 136–142.

Kopera HC, Larson PA, Moldovan JB, Richardson SR, Liu Y, Moran J V. 2016. Line-1 cultured cell retrotransposition assay. In Methods in Molecular Biology, Vol. 1400 of, pp. 139–156, Humana Press Inc.

Kubo S, Seleme M d. C, Soifer HS, Perez JLG, Moran JV, Kazazian HH, Kasahara N. 2006. L1 retrotransposition in nondividing and primary human somatic cells. Proc Natl Acad Sci 103: 8036– 8041.

Kumaki Y, Oda M, Okano M. 2008. QUMA: quantification tool for methylation analysis. Nucleic Acids Res 36: W170–5.

Kuramochi-Miyagawa S, Watanabe T, Gotoh K, Totoki Y, Toyoda A, Ikawa M, Asada N, Kojima K, Yamaguchi Y, Ijiri TW, et al. 2008. DNA methylation of retrotransposon genes is regulated by Piwi family members MILI and MIWI2 in murine fetal testes. Genes Dev 22: 908–917.

Lanciano S, Cristofari G. 2020. Measuring and interpreting transposable element expression. Nat Rev Genet 21: 721–736.

Lander ES, Linton LM, Birren B, Nusbaum C, Zody MC, Baldwin J, Devon K, Dewar K, Doyle M, FitzHugh W, et al. 2001. Initial sequencing and analysis of the human genome. Nature 409: 860–921.

Lee S-H, Cho S-Y, Shannon MF, Fan J, Rangasamy D. 2010. The Impact of CpG Island on Defining Transcriptional Activation of the Mouse L1 Retrotransposable Elements ed. I.K. Jordan. PLoS One 5: e11353.

Li H. 2018. Minimap2: Pairwise alignment for nucleotide sequences. Bioinformatics 34: 3094–3100.

Li H. 2011. Tabix: Fast retrieval of sequence features from generic TAB-delimited files. Bioinformatics 27: 718–719.

Li H, Handsaker B, Wysoker A, Fennell T, Ruan J, Homer N, Marth G, Abecasis G, Durbin R. 2009. The Sequence Alignment/Map format and SAMtools. Bioinformatics 25: 2078–2079.

Liu N, Lee CH, Swigut T, Grow E, Gu B, Bassik MC, Wysocka J. 2018. Selective silencing of euchromatic L1s revealed by genome-wide screens for L1 regulators. Nature 553: 228–232.

Loeb DD, Padgett RW, Hardies SC, Shehee WR, Comer MB, Edgell MH, Hutchison CA. 1986. The sequence of a large L1Md element reveals a tandemly repeated 5’ end and several features found in retrotransposons. Mol Cell Biol 6: 168–182.

Luan DD, Korman MH, Jakubczak JL, Eickbush TH. 1993. Reverse transcription of R2Bm RNA is primed by a nick at the chromosomal target site: A mechanism for non-LTR retrotransposition. Cell 72: 595– 605.

Macia A, Widmann TJ, Heras SR, Ayllon V, Sanchez L, Benkaddour-Boumzaouad M, Muñoz-Lopez M, Rubio A, Amador-Cubero S, Blanco-Jimenez E, et al. 2017. Engineered LINE-1 retrotransposition in nondividing human neurons. Genome Res 27: 335–348.

MacLennan M, García-Cañadas M, Reichmann J, Khazina E, Wagner G, Playfoot CJ, Salvador-Palomeque C, Mann AR, Peressini P, Sanchez L, et al. 2017. Mobilization of LINE-1 retrotransposons is restricted by Tex19.1 in mouse embryonic stem cells. Elife 6: e26152.

Magoč T, Salzberg SL. 2011. FLASH: Fast length adjustment of short reads to improve genome assemblies. Bioinformatics 27: 2957–2963.

Martin SL, Bushman FD. 2001. Nucleic acid chaperone activity of the ORF1 protein from the mouse LINE-1 retrotransposon. Mol Cell Biol 21: 467–75.

Mathias S, Scott A, Kazazian HH, Boeke J, Gabriel A. 1991. Reverse transcriptase encoded by a human transposable element. Science 254: 1808–1810.

Mears ML, Hutchison CA. 2001. The evolution of modern lineages of mouse L1 elements. J Mol Evol 52: 51–62.

Mita P, Sun X, Fenyö D, Kahler DJ, Li D, Agmon N, Wudzinska A, Keegan S, Bader JS, Yun C, et al. 2020. BRCA1 and S phase DNA repair pathways restrict LINE-1 retrotransposition in human cells. Nat Struct Mol Biol 27: 179–191.

Mita P, Wudzinska A, Sun X, Andrade J, Nayak S, Kahler DJ, Badri S, LaCava J, Ueberheide B, Yun CY, et al. 2018. LINE-1 protein localization and functional dynamics during the cell cycle. Elife 7: e30058.

Molaro A, Falciatori I, Hodges E, Aravin AA, Marran K, Rafii S, McCombie WR, Smith AD, Hannon GJ. 2014. Two waves of de novo methylation during mouse germ cell development. Genes Dev 28: 1544– 9.

Moran JV, DeBerardinis RJ, Kazazian HH. 1999. Exon shuffling by L1 retrotransposition. Science 283: 1530–4.

Moran JV, Holmes SE, Naas TP, DeBerardinis RJ, Boeke JD, Kazazian HH. 1996. High frequency retrotransposition in cultured mammalian cells. Cell 87: 917–27.

Mottez E, Rogan PK, Manuelidis L. 1986. Conservation in the 5’ region of the long interspersed mouse L1 repeat: implications of comparative sequence analysis. Nucleic Acids Res 14: 3119–3136.

Naas TP, DeBerardinis RJ, Moran JV, Ostertag EM, Kingsmore SF, Seldin MF, Hayashizaki Y, Martin SL, Kazazian HH. 1998. An actively retrotransposing, novel subfamily of mouse L1 elements. EMBO J 17: 590–597.

Nellåker C, Keane TM, Yalcin B, Wong K, Agam A, Belgard TG, Flint J, Adams DJ, Frankel WN, Ponting CP. 2012. The genomic landscape shaped by selection on transposable elements across 18 mouse strains. Genome Biol 13: R45.

Nguyen THM, Carreira PE, Sanchez-Luque FJ, Schauer SN, Fagg AC, Richardson SR, Davies CM, Jesuadian JS, Kempen MJHC, Troskie RL, et al. 2018. L1 Retrotransposon Heterogeneity in Ovarian Tumor Cell Evolution. Cell Rep 23: 3730–3740.

Okano M, Xie S, Li E. 1998. Cloning and characterization of a family of novel mammalian DNA (cytosine-5) methyltransferases. Nat Genet 19: 219–220.

Ostertag EM, Kazazian HH. 2001. Twin priming: a proposed mechanism for the creation of inversions in L1 retrotransposition. Genome Res 11: 2059–65.

Paterson AL, Weaver JMJ, Eldridge MD, Tavaré S, Fitzgerald RC, Edwards PAW. 2015. Mobile element insertions are frequent in oesophageal adenocarcinomas and can mislead paired-end sequencing analysis. BMC Genomics 16: 1–14.

Penzkofer T, Jäger M, Figlerowicz M, Badge R, Mundlos S, Robinson PN, Zemojtel T. 2017. L1Base 2: More retrotransposition-active LINE-1s, more mammalian genomes. Nucleic Acids Res 45: D68–D73.

Philippe C, Vargas-Landin DB, Doucet AJ, Van Essen D, Vera-Otarola J, Kuciak M, Corbin A, Nigumann P, Cristofari G. 2016. Activation of individual L1 retrotransposon instances is restricted to cell-type dependent permissive loci. Elife 5: e13926.

Pickeral OK, Makałowski W, Boguski MS, Boeke JD. 2000. Frequent human genomic DNA transduction driven by LINE-1 retrotransposition. Genome Res 10: 411–5.

Pitkänen E, Cajuso T, Katainen R, Kaasinen E, Välimäki N, Palin K, Taipale J, Aaltonen LA, Kilpivaara O. 2014. Frequent L1 retrotranspositions originating from TTC28 in colorectal cancer. Oncotarget 5: 853–859.

Popp C, Dean W, Feng S, Cokus SJ, Andrews S, Pellegrini M, Jacobsen SE, Reik W. 2010. Genome-wide erasure of DNA methylation in mouse primordial germ cells is affected by AID deficiency. Nature 463: 1101–1105.

Richardson SR, Gerdes P, Gerhardt DJ, Sanchez-Luque FJ, Bodea GO, Muñoz-Lopez M, Jesuadian JS, Kempen MJHC, Carreira PE, Jeddeloh JA, et al. 2017. Heritable L1 retrotransposition in the mouse primordial germline and early embryo. Genome Res 27: 1395–1405.

Saitou M, Kagiwada S, Kurimoto K. 2012. Epigenetic reprogramming in mouse pre-implantation development and primordial germ cells. Development 139: 15–31.

Salvador-Palomeque C, Sanchez-Luque FJ, Fortuna PRJ, Ewing AD, Wolvetang EJ, Richardson SR, Faulkner GJ. 2019. Dynamic Methylation of an L1 Transduction Family during Reprogramming and Neurodifferentiation. Mol Cell Biol 39: e00499–18.

Sanchez-Luque FJ, Kempen MJHC, Gerdes P, Vargas-Landin DB, Richardson SR, Troskie RL, Jesuadian JS, Cheetham SW, Carreira PE, Salvador-Palomeque C, et al. 2019. LINE-1 Evasion of Epigenetic Repression in Humans. Mol Cell 75: 590–604.e12.

Saxton JA, Martin SL. 1998. Recombination between subtypes creates a mosaic lineage of LINE-1 that is expressed and actively retrotransposing in the mouse genome. J Mol Biol 280: 611–622.

Schauer SN, Carreira PE, Shukla R, Gerhardt DJ, Gerdes P, Sanchez-Luque FJ, Nicoli P, Kindlova M, Ghisletti S, Dos Santos AD, et al. 2018. L1 retrotransposition is a common feature of mammalian hepatocarcinogenesis. Genome Res 28: 639–653.

Schichman SA, Adey NB, Edgell MH, Hutchison CA. 1993. L1 A-monomer tandem arrays have expanded during the course of mouse L1 evolution. Mol Biol Evol 10: 552–570.

Schöpp T, Zoch A, Berrens RV, Auchynnikava T, Kabayama Y, Vasiliauskaitė L, Rappsilber J, Allshire RC, O’Carroll D. 2020. TEX15 is an essential executor of MIWI2-directed transposon DNA methylation and silencing. Nat Commun 11: 1–8.

Scott AF, Schmeckpeper B, Abdelrazik M, Comey C, O’Hara B, Rossiter J, Cooley T, Heath P, Smith K, Margolet L. 1987. Origin of the human L1 elements: proposed progenitor genes deduced from a consensus DNA sequence. Genomics 1: 113–25.

Scott EC, Gardner EJ, Masood A, Chuang NT, Vertino PM, Devine SE. 2016. A hot L1 retrotransposon evades somatic repression and initiates human colorectal cancer. Genome Res 26: 745–755.

Seisenberger S, Andrews S, Krueger F, Arand J, Walter J, Santos F, Popp C, Thienpont B, Dean W, Reik W, et al. 2012. The dynamics of genome-wide DNA methylation reprogramming in mouse primordial germ cells. Mol Cell 48: 849–62.

Seki Y, Hayashi K, Itoh K, Mizugaki M, Saitou M, Matsui Y. 2005. Extensive and orderly reprogramming of genome-wide chromatin modifications associated with specification and early development of germ cells in mice. Dev Biol 278: 440–458.

Senft AD, Macfarlan TS. 2021. Transposable elements shape the evolution of mammalian development. Nat Rev Genet 22: 691–711.

Severynse DM, Hutchison CA, Edgell MH. 1991. Identification of transcriptional regulatory activity within the 5′ A-type monomer sequence of the mouse LINE-1 retroposon. Mamm Genome 2: 41–50.

Shehee WR, Chao SF, Loeb DD, Comer MB, Hutchison CA, Edgell MH. 1987. Determination of a functional ancestral sequence and definition of the 5′ end of A-type mouse L1 elements. J Mol Biol 196: 757–767.

Silva J, Barrandon O, Nichols J, Kawaguchi J, Theunissen TW, Smith A. 2008. Promotion of Reprogramming to Ground State Pluripotency by Signal Inhibition ed. M.A. Goodell. PLoS Biol 6: e253.

Simpson JT, Workman RE, Zuzarte PC, David M, Dursi LJ, Timp W. 2017. Detecting DNA cytosine methylation using nanopore sequencing. Nat Methods 14: 407–410.

Skowronski J, Fanning TG, Singer MF. 1988. Unit-length LINE-1 transcripts in human teratocarcinoma cells. Mol Cell Biol 8: 1385–97.

Slater GSC, Birney E. 2005. Automated generation of heuristics for biological sequence comparison. BMC Bioinformatics 6: 31.

Smith AG, Heath JK, Donaldson DD, Wong GG, Moreau J, Stahl M, Rogers D. 1988. Inhibition of pluripotential embryonic stem cell differentiation by purified polypeptides. Nature 336: 688–690.

Smith ZD, Chan MM, Mikkelsen TS, Gu H, Gnirke A, Regev A, Meissner A. 2012. A unique regulatory phase of DNA methylation in the early mammalian embryo. Nature 484: 339–344.

Sookdeo A, Hepp CM, McClure MA, Boissinot S. 2013. Revisiting the evolution of mouse LINE-1 in the genomic era. Mob DNA 4: 3.

Symer DE, Connelly C, Szak ST, Caputo EM, Cost GJ, Parmigiani G, Boeke JD, Moran JV, Kazazian HH, Mehdi SQ, et al. 2002. Human L1 retrotransposition is associated with genetic instability in vivo. Cell 110: 327–38.

Tachibana M, Nozaki M, Takeda N, Shinkai Y. 2007. Functional dynamics of H3K9 methylation during meiotic prophase progression. EMBO J 26: 3346–59.

Tan X, Xu X, Elkenani M, Smorag L, Zechner U, Nolte J, Engel W, Pantakani DVK. 2013. Zfp819, a novel KRAB-zinc finger protein, interacts with KAP1 and functions in genomic integrity maintenance of mouse embryonic stem cells. Stem Cell Res 11: 1045–1059.

Taylor MS, LaCava J, Mita P, Molloy KR, Huang CRL, Li D, Adney EM, Jiang H, Burns KH, Chait BT, et al. 2013. Affinity Proteomics Reveals Human Host Factors Implicated in Discrete Stages of LINE-1 Retrotransposition. Cell 155: 1034–1048.

Tristán-Ramos P, Rubio-Roldan A, Peris G, Sánchez L, Amador-Cubero S, Viollet S, Cristofari G, Heras SR. 2020. The tumor suppressor microRNA let-7 inhibits human LINE-1 retrotransposition. Nat Commun 11: 5712.

Tubio JMC, Li Y, Ju YS, Martincorena I, Cooke SL, Tojo M, Gundem G, Pipinikas CP, Zamora J, Raine K, et al. 2014. Extensive transduction of nonrepetitive DNA mediated by L1 retrotransposition in cancer genomes. Science 345: 1251343–1251343.

Voliva CF, Jahn CL, Comer MB, Hutchison CA, Edgell MH. 1983. The L1Md long interspersed repeat family in the mouse: Almost all examples are truncated at one end. Nucleic Acids Res 11: 8847–8859.

Walter M, Teissandier A, Pérez-Palacios R, Bourc’his D. 2016. An epigenetic switch ensures transposon repression upon dynamic loss of DNA methylation in embryonic stem cells. Elife 5: e11418.

Watanabe T, Totoki Y, Toyoda A, Kaneda M, Kuramochi-Miyagawa S, Obata Y, Chiba H, Kohara Y, Kono T, Nakano T, et al. 2008. Endogenous siRNAs from naturally formed dsRNAs regulate transcripts in mouse oocytes. Nature 453: 539–543.

Waterston RH, Lindblad-Toh K, Birney E, Rogers J, Abril JF, Agarwal P, Agarwala R, Ainscough R, Alexandersson M, An P, et al. 2002. Initial sequencing and comparative analysis of the mouse genome. Nature 420: 520–62.

Wei W, Morrish TA, Alisch RS, Moran JV. 2000. A transient assay reveals that cultured human cells can accommodate multiple LINE-1 retrotransposition events. Anal Biochem 284: 435–438.

Williams RL, Hilton DJ, Pease S, Willson TA, Stewart CL, Gearing DP, Wagner EF, Metcalf D, Nicola NA, Gough NM. 1988. Myeloid leukaemia inhibitory factor maintains the developmental potential of embryonic stem cells. Nature 336: 684–687.

Wincker P, Jubier-Maurin V, Roizés G. 1987. Unrelated sequences at the 5′ end of mouse LINE-1 repeated elements define two distinct sub-families. Nucleic Acids Res 15: 8593–8606.

Xing J, Wang H, Belancio VP, Cordaux R, Deininger PL, Batzer MA. 2006. Emergence of primate genes by retrotransposon-mediated sequence transduction. Proc Natl Acad Sci U S A 103: 17608–13.

Zhou M, Smith AD. 2019. Subtype classification and functional annotation of L1Md retrotransposon promoters. Mob DNA 10: 14.

Zingler N, Willhoeft U, Brose H-P, Schoder V, Jahns T, Hanschmann K-MO, Morrish TA, Löwer J, Schumann GG. 2005. Analysis of 5’ junctions of human LINE-1 and Alu retrotransposons suggests an alternative model for 5’-end attachment requiring microhomology-mediated end-joining. Genome Res 15: 780–9.

Zoch A, Auchynnikava T, Berrens RV, Kabayama Y, Schöpp T, Heep M, Vasiliauskaitė L, Pérez-Rico YA, Cook AG, Shkumatava A, et al. 2020. SPOCD1 is an essential executor of piRNA-directed de novo DNA methylation. Nature 584: 635–639.

