## Supplemental Figures for "Mouse L1s fade with age: a methylation-enforced mechanism for attenuation of L1 retrotransposition potential"

### Supplemental Figure S1

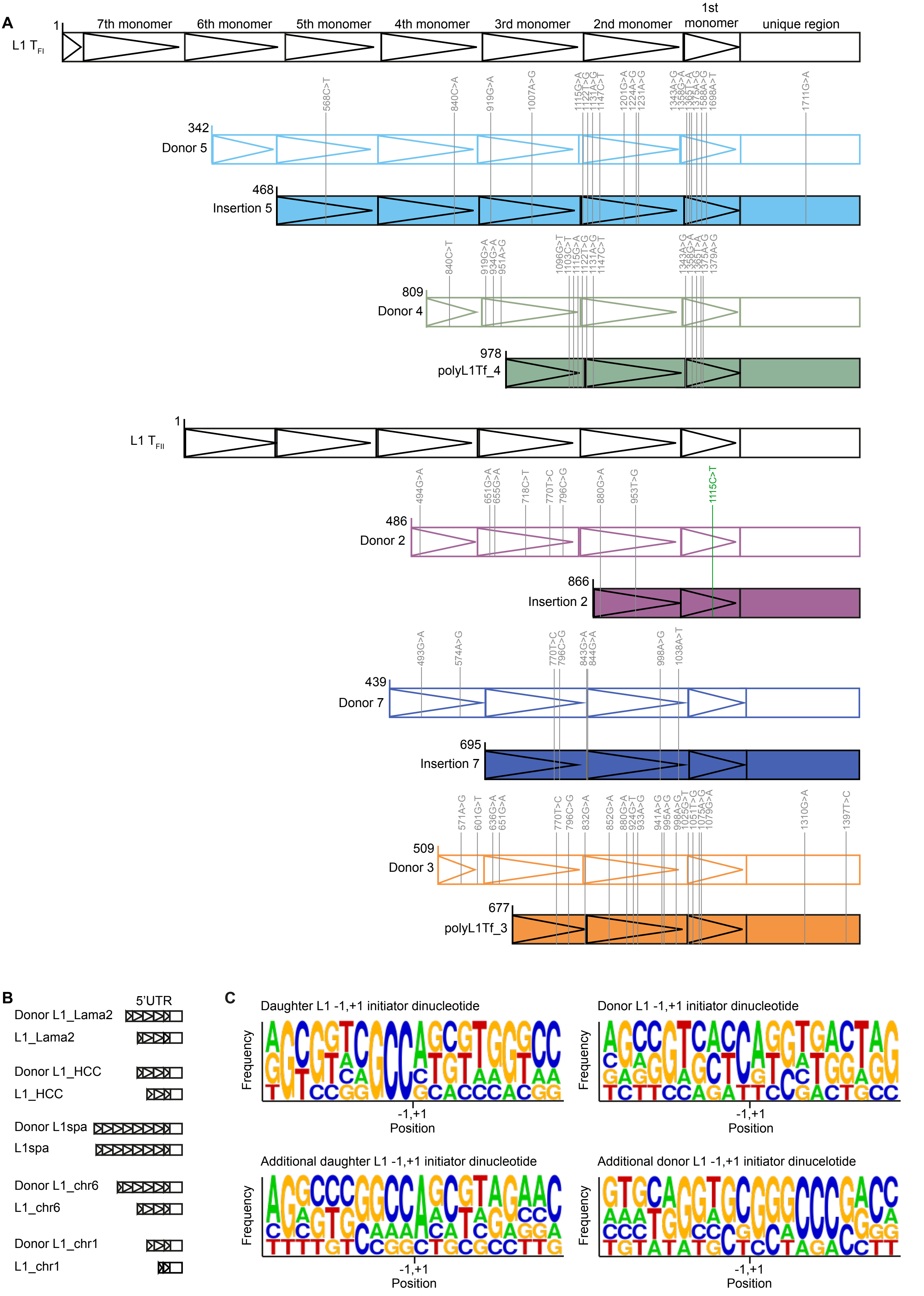

**Supplemental Figure S1. L1 donor/daughter promoter sequence comparison.**

(*A*) Sequence comparison of L1 $T_{FI}$ and $T_{FII}$ donor/daughter pair promoters. Nucleotide substitutions compared to the consensus sequence are annotated in grey. Nucleotide substitution in green represents mutation within extended YY1 binding motif. Triangles within 5′ UTR represent monomer units.

(*B*) Cartoons for promoters of 5 additional L1 donor/daughter pairs identified in the literature and our analysis.

(*C*) Sequence logos (Crooks et al. 2004) of putative transcription initiation start sites for donor and daughter L1s. Sequence represents -1,+1 transcription initiator dinucleotide in the center ± 9 nucleotides upstream and downstream. +1 indicates first nucleotide of L1 sequence which corresponds to second nucleotide in transcription initiator dinucleotide.

### Supplemental Figure S2

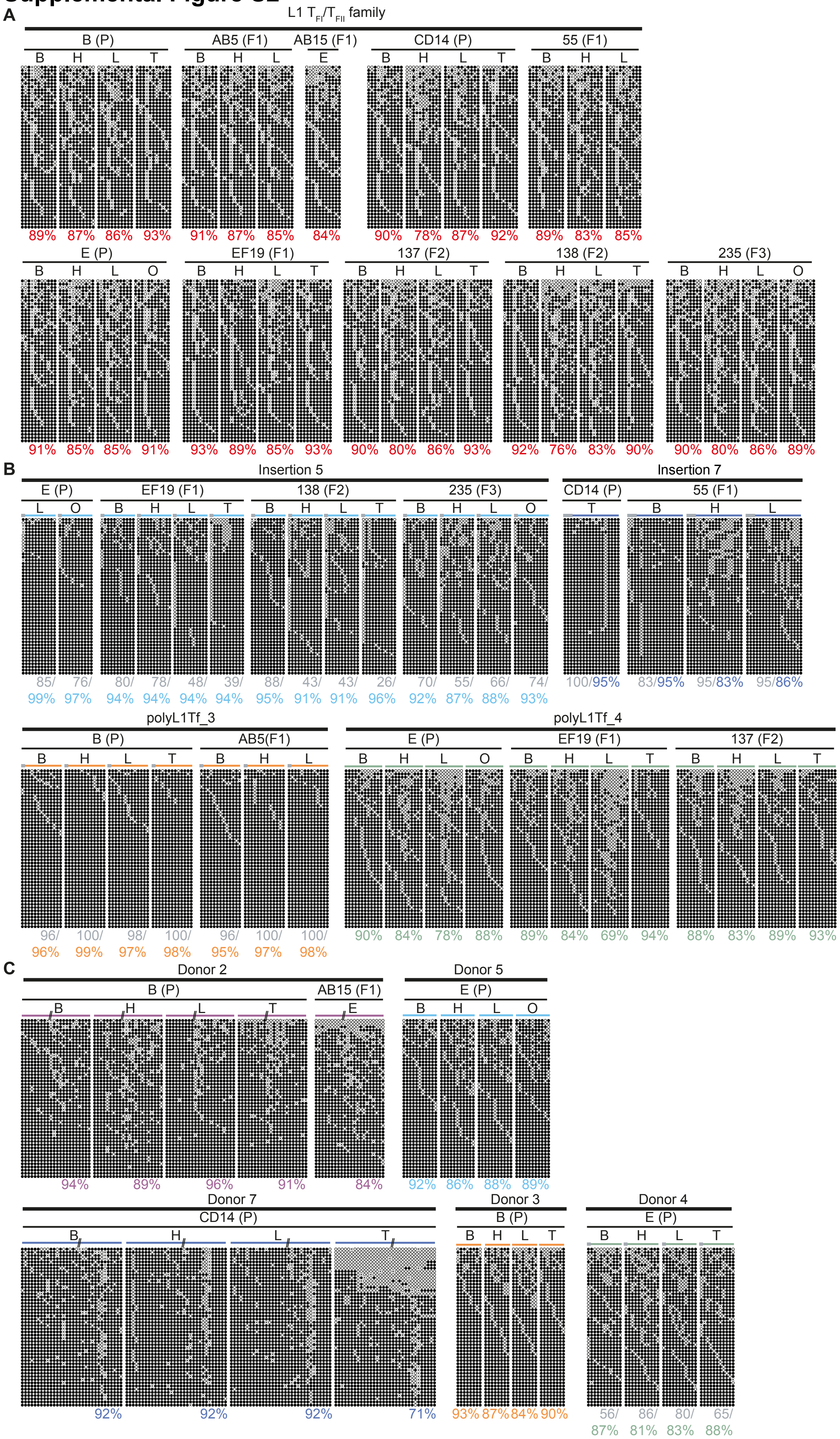

**Supplemental Figure S2. Methylation of L1 $T_{FI}$/ $T_{FII}$ family and L1 donor/daughter pairs in mouse tissues.**

(*A*) Genome-wide methylation level of CpG dinucleotides of L1 $T_{FI}$/ $T_{FII}$ promoter sequence is shown. Displayed are 50 non-identical sequences extracted at random from a much larger pool of available Illumina reads. Each cartoon panel corresponds to an amplicon (black circle, methylated CpG; white circle, unmethylated CpG; ×, mutated CpG). The overall percentage of methylated CpG dinucleotides is indicated below each cartoon.

(*B*) As for (*A*) except for methylation level of CpG dinucleotides of L1 daughter insertions is shown. Colored line above cartoon represents amplicon (grey = genomic sequence, colored = L1 sequence). Grey letters indicate overall percentage of methylated CpG dinucleotides in genomic sequence. Colored letters indicate methylation of CpG dinucleotides in L1 sequence.

(*C*) As for (*A*) and (*B*) but for donor L1s. The promoters of Donor 2 and Donor 7 were not completely sequenced and missing the center monomer as indicated by black lines in colored line above methylation cartoons.

### Supplemental Figure S3

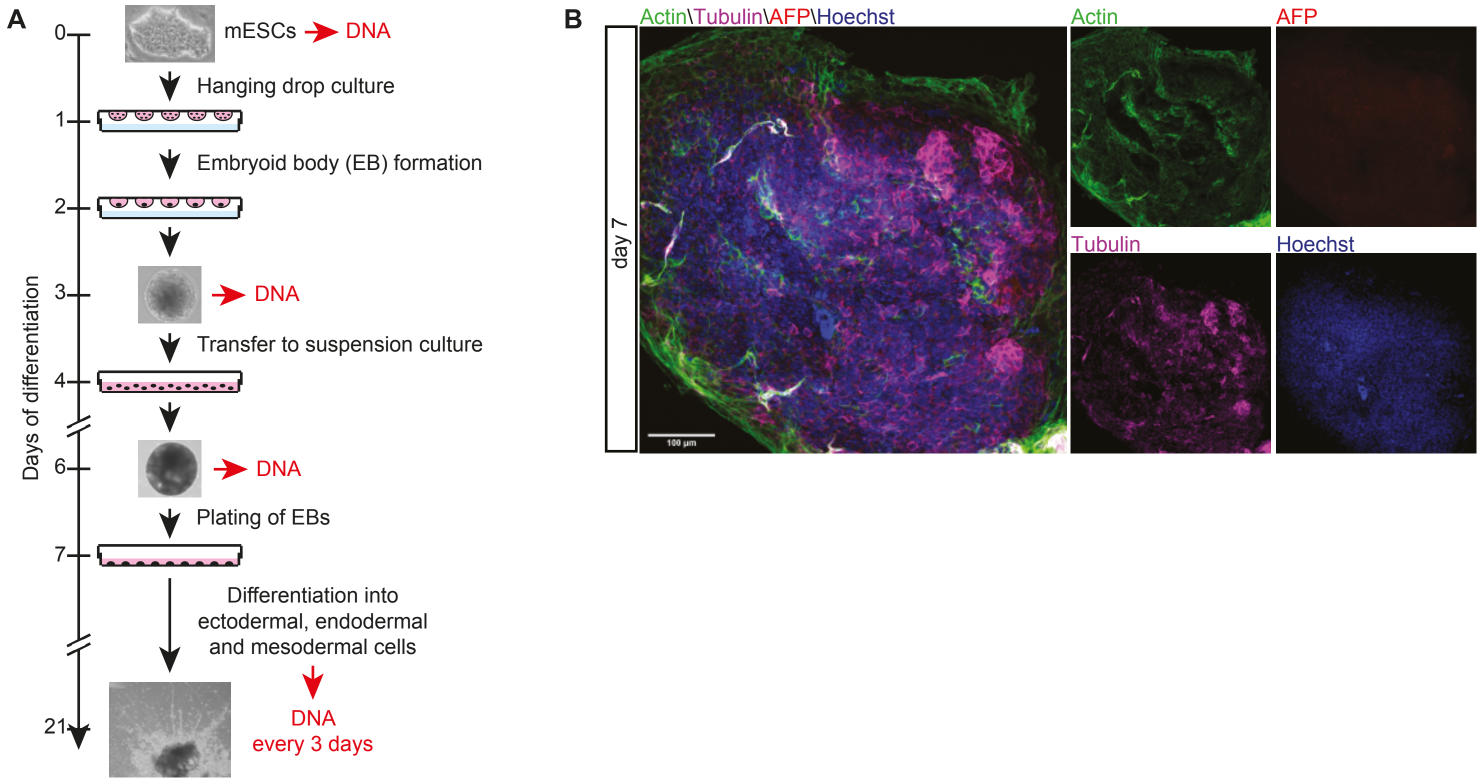

**Supplemental Figure S3. Differentiation of E14 mESCs.**

(*A*) Schematic of mESC differentiation by hanging drop culture. Differentiation of mESCs was induced by culturing mESCs in hanging drops to induce embryoid body (EB) formation. On day three EBs were transferred to low-attachment tissue culture dishes. After 6 days, EBs were plated and differentiated for 14 days. On day 21, differentiated cells were fixed and stained for markers of ectodermal, endodermal and mesodermal cells. Cells were collected every 3 days during differentiation and DNA for methylation analysis was extracted.

(*B*) Immunofluorescence image of mesodermal (Actin), endodermal (AFP) and ectodermal (Tubulin) lineage markers in differentiated E14 mESCs on d7 (one day after plating EBs). Nuclei were stained with Hoechst (blue). Scale bar, 100 µm.

### Supplemental Figure S4

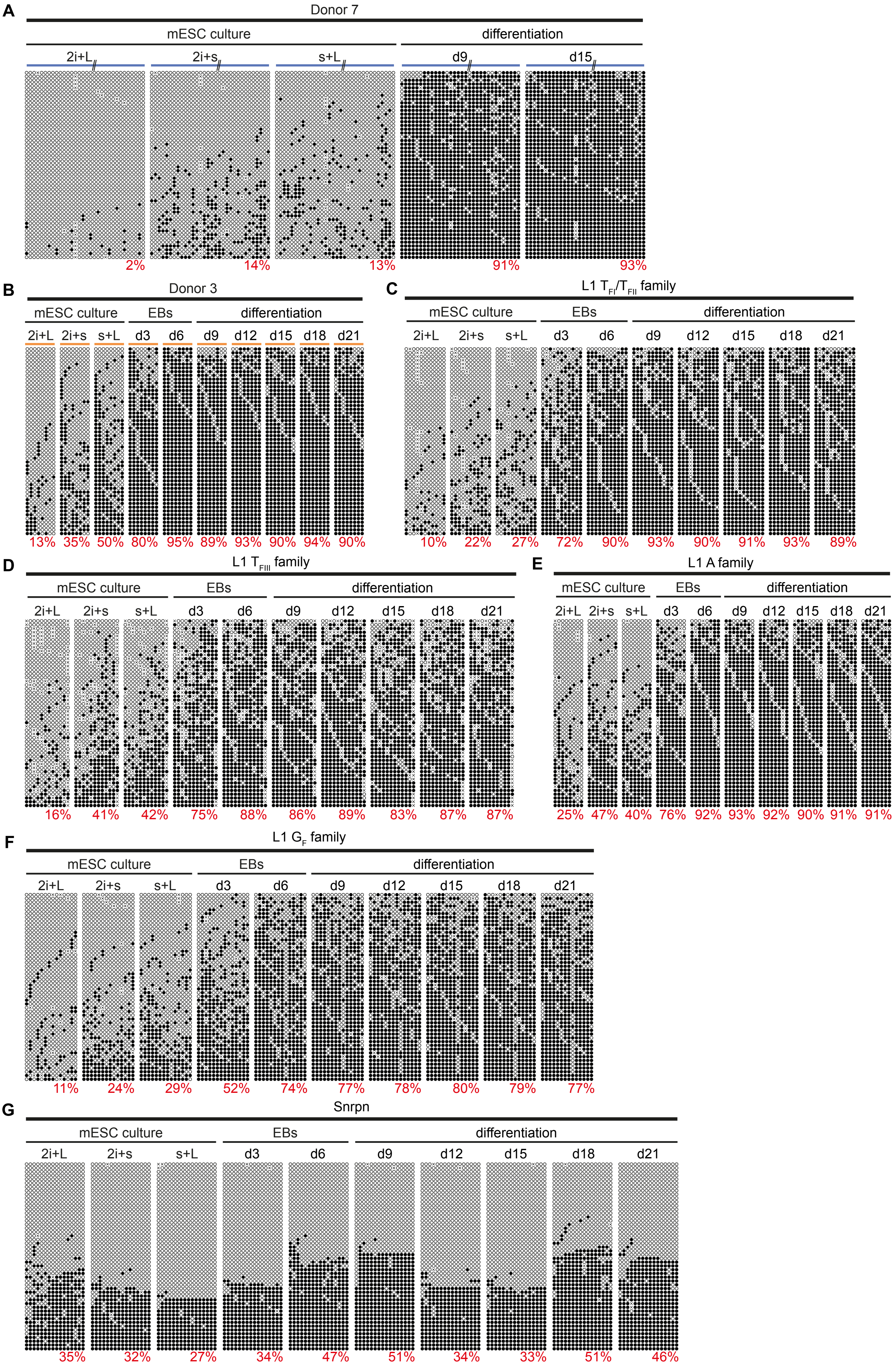

**Supplemental Figure S4. Dynamic methylation during mESC differentiation.**

(*A*) Methylation of Donor 7 promoter sequence shown in the mESCs cultured in three different conditions (2i+L = 2i+LIF, 2i+s = 2i+serum, s+L = serum+LIF) and on differentiation day 9 (d9) and day 15 (d15). Displayed are 50 non-identical sequences extracted at random from a much larger pool of available Illumina reads. Each cartoon panel corresponds to an amplicon (black circle, methylated CpG; white circle, unmethylated CpG; ×, mutated CpG). Colored line above each cartoon represents amplicon (grey = genomic sequence, colored = L1 sequence). The overall percentage of methylated CpG dinucleotides is indicated below each cartoon. Grey letters indicate methylation of CpG dinucleotides in genomic sequence. Colored letters indicate methylation of CpG dinucleotides in L1 sequence. The promoter of Donor 7 was not completely sequenced as indicated by black lines in colored line above methylation cartoons.

(*B-F*) As per (*A*) but for Donor 3 (*B*), L1 $T_{FI}$/$T_{FII}$ family (*C*), L1 $T_{FIII}$ family (*D*), L1 A family (*E*), L1 $G_F$ family (*F*). Primers for L1 subfamilies are within the L1 promoter sequence. Shown is methylation in three different mESC culture conditions, during EB culture and during differentiation.

(*G*) As per above but for the imprinted gene *Snrpn*.

### Supplemental Figure S5

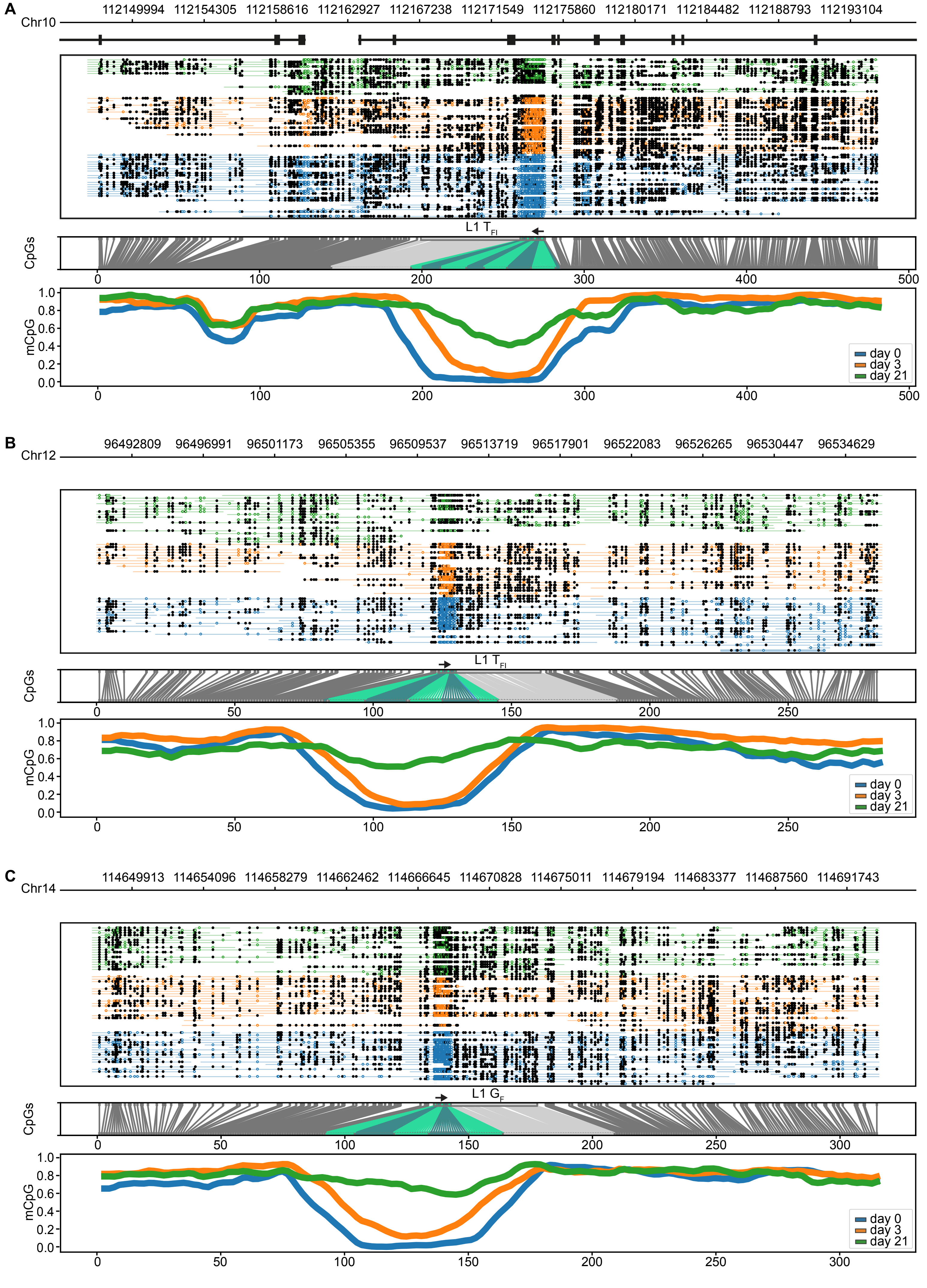

**Supplemental Figure S5. ONT methylation profiles for L1 somatic methylation "escapee" loci.**

(*A*) Methylation of an L1 $T_{FI}$ element on chromosome 10 and surrounding locus. From *top* to *bottom* this figure shows i) the genomic position of the L1 in an intron on chromosome 10, including 20 kbp up- and downstream of the L1, ii) a diagram showing methylated (filled black circles) and unmethylated (unfilled colored circles) CpGs and read (colored lines) coverage per sample, iii) a diagram displaying the correspondence between genome space and CpG space, CpGs belonging to full-length L1 are annotated in light and dark green (promoter monomers) and light grey (ORFs and 3′ UTR), iv) the fraction of methylated CpGs for three differentiation time points (d0, d3, d21) in CpG space.

(*B*) As for (*A*) except for an intergenic L1 $T_{FI}$ element on chromosome 12.

(*C*) As for (*A*) except for an intergenic L1 $G_F$ element on chromosome 14.

### Supplemental Figure S6

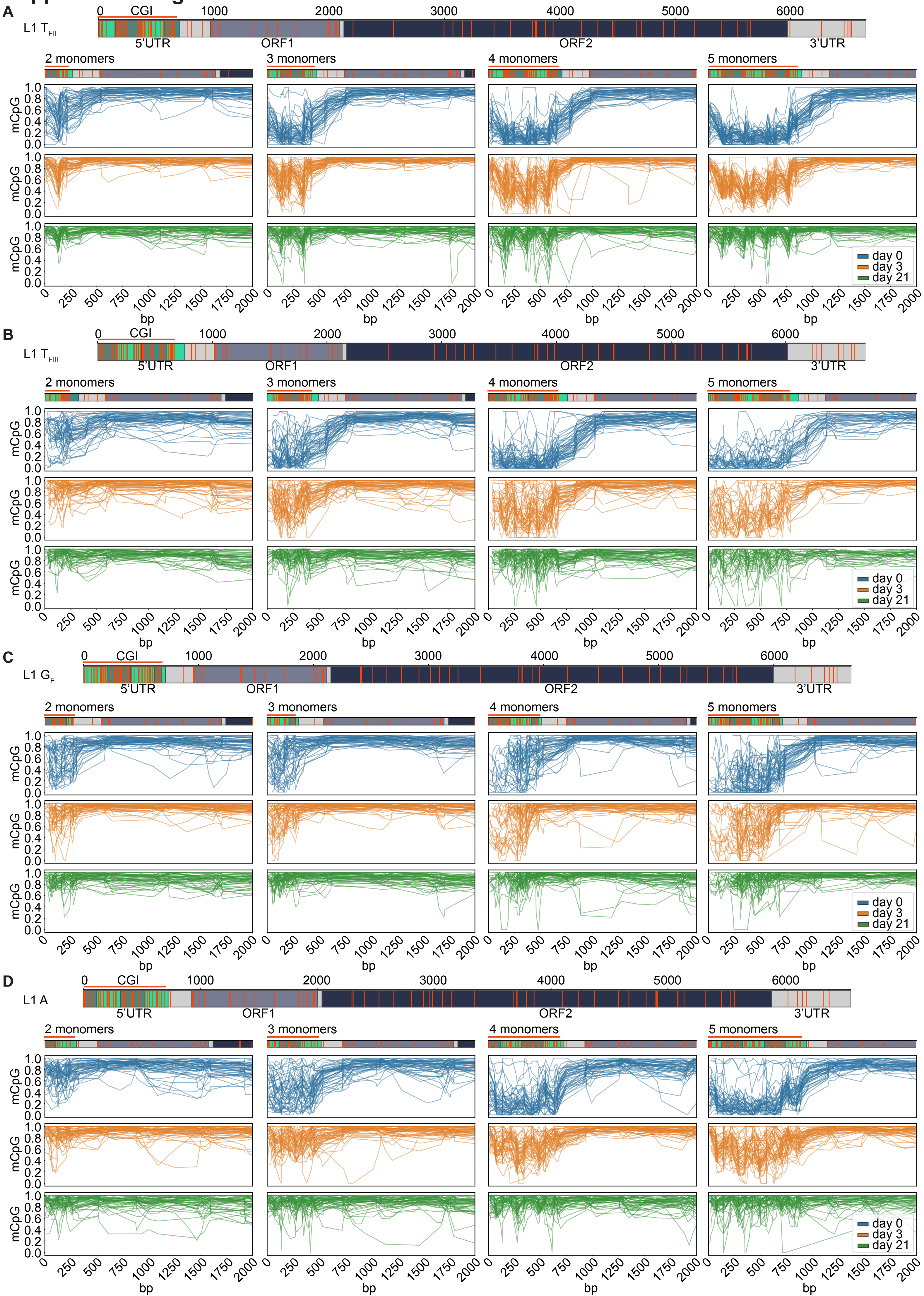

**Supplemental Figure S6. Composite methylation profiles of L1 subfamily promoters during differentiation.**

(*A*) *Top*: Annotated full-length L1 $T_{FII}$ consensus showing the monomer units in green, unique region in light grey, ORF1 in dark grey, ORF2 in dark green, and 3′ UTR in light grey. CpG dinucleotides throughout the whole element are displayed as orange strokes. The promoter CpG island (CGI) is indicated as an orange line. Number of bp are shown above the element.
*Bottom*: Data is shown for L1 $T_{FII}$ promoters containing 2, 3, 4 and 5 monomers at three time points of differentiation: d0 (undifferentiated mESCs in serum+LIF), d3 (EBs on day 3 of differentiation) and on d21 (completely differentiated cells). Each graph displays up to 50 methylation profiles. Annotated consensus sequences as per (*top*) are shown at *top* including CpG positions.

(*B*) As for (*A*) except for L1 $T_{FIII}$ promoters.

(*C*) As for (*A*) except for L1 $G_F$ promoters.

(*D*) As for (*A*) except for L1 A promoters.

### Supplemental Figure S7

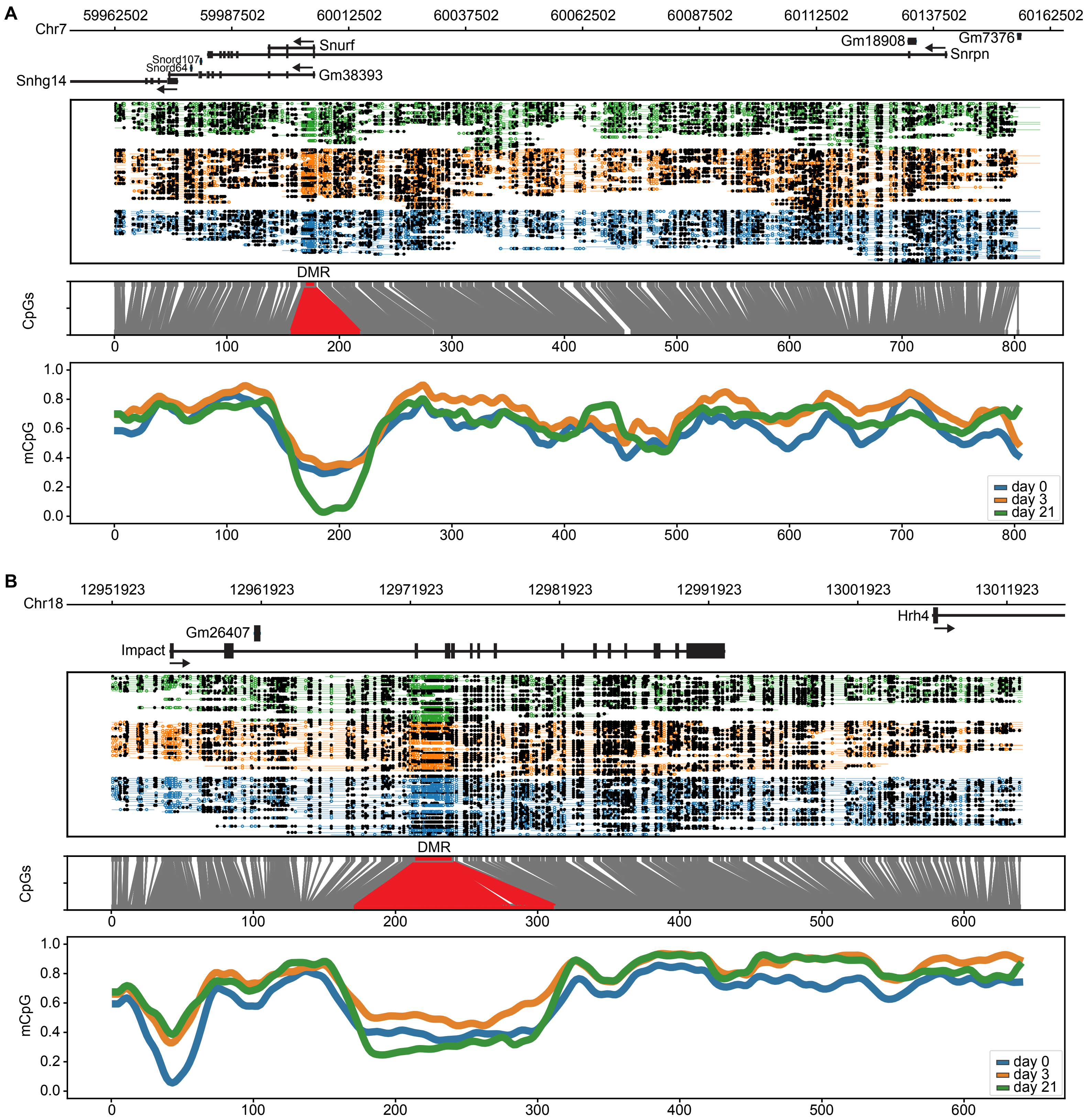

**Supplemental Figure S7. ONT methylation profiles of imprinted genes *Snrpn* and *Impact*.**

(*A*) Methylation of the *Snrpn* gene and surrounding locus. From *top* to *bottom* this figure shows i) the genomic position of *Snrpn* on chromosome 7, including 20 kbp up- and downstream of *Snrpn*, ii) a diagram showing methylated (filled black circles) and unmethylated (unfilled colored circles) CpGs and read (colored lines) coverage per sample, (iii) a diagram displaying the correspondence between genome space and CpG space, CpGs belonging to the *Snrpn* DMR are annotated in red, iv) the fraction of methylated CpGs for three differentiation time points (d0, d3, d21) in CpG space.

(*B*) As for (*A*) except for the imprinted *Impact* gene.
